# A Personalized Approach to Student Mental Health: Real-Time EEG Biofeedback for Stress and Focus Management

**DOI:** 10.64898/2026.10.20.759906

**Authors:** Ankit Kumar Jha, Ali Kalsekar, Mohammed Salah Chogle

**Affiliations:** Automation and Robotics Engineering, Vivekanand Education Society’s Institute of Technology, Chembur

**Keywords:** EEG, stress-state estimation, biofeedback, Fast Fourier Transform, Beta/Alpha ratio, personalized thresholding, brain–computer interface

## Abstract

Real-time monitoring of student stress and focus requires low-cost EEG systems, but fixed spectral thresholds may not transfer reliably across users because EEG amplitude and band power vary between individuals and recording conditions. This study investigates a subject-specific thresholding approach using a two-channel Brain BioAmp acquisition setup (FP1 and FP2) and an ESP32-based FFT pipeline. Beta/Alpha and Theta/Alpha ratios are calculated from spectral band estimates and compared with thresholds derived from Baseline, Focused, and Stressed task-condition recordings. In the reported pilot results, the mean Beta/Alpha ratio increased from 2.76 during Baseline to 2.80 during Focused and 2.88 during Stressed conditions. The Theta/Alpha ratio was lowest during Focused (0.95) and highest during Stressed (1.04), compared with 0.99 during Baseline. These are descriptive observations; classification accuracy, statistical significance, and generalization across participants have not been established. The signal-processing calculations and plotted frequency ranges should be checked against the original implementation before these values are treated as final.

## 1. Introduction

Electroencephalography (EEG) is used to observe rhythmic electrical activity associated with neuronal populations. EEG activity is commonly described using frequency bands such as delta, theta, alpha, beta, and gamma, which provide useful features for studying changes in brain activity. Interpretation of these features requires care because EEG signals are affected by individual differences, electrode placement, recording conditions, and physiological artifacts [1]–[3].

This work investigates a low-cost, two-channel frontal EEG pipeline for comparing spectral features across Baseline, Focused, and Stressed task conditions. The intended contributions are: (i) an acquisition and FFT-processing pipeline implemented with an ESP32; (ii) a subject-specific thresholding procedure based on Beta/Alpha and Theta/Alpha ratios; and (iii) a pilot comparison of these features across three task conditions. The study is a feasibility investigation and does not establish clinical or general-purpose stress-detection performance.

## 2. Related Work and Research Gap

Prior EEG research has examined spectral characteristics and signal processing [1]–[3], brain-signal control [4], classification methods [5], biometrics [6], emotion-related features [7], and brain–computer interfaces [8], [9]. Surveys have also discussed EEG analysis and machine-learning approaches to EEG processing [10], [11]. Research on wearable systems and student-focused stress monitoring provides additional context for applying physiological sensing in educational settings [12]–[14].

EEG features vary across individuals and recording conditions, so a threshold selected from one person or session may not generalize to another. This study therefore investigates subject-specific midpoint thresholds in a low-cost frontal EEG setup. The contribution is an implementation-oriented feasibility evaluation; comparative accuracy and generalization remain to be established.

## 3. Methodology

**Fig. 1.**
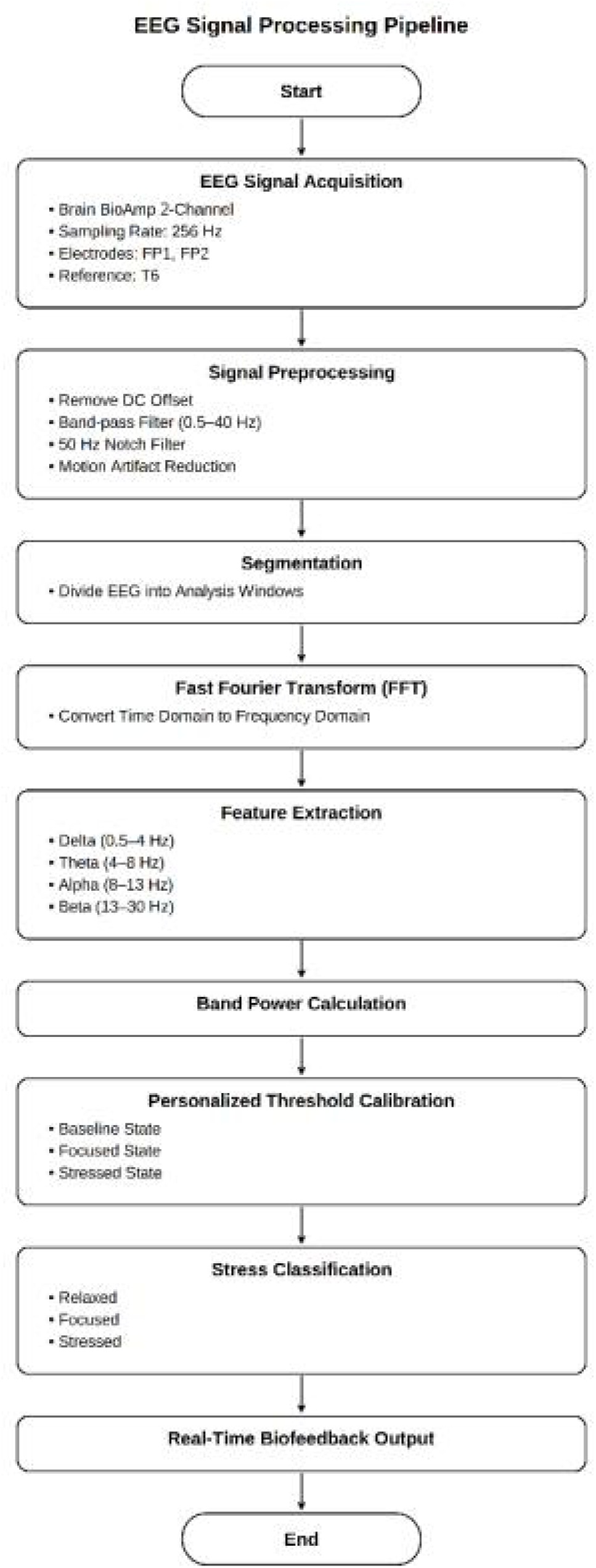
EEG signal-processing pipeline.

The processing workflow consists of EEG acquisition, preprocessing, windowing, FFT analysis, spectral feature calculation, feature-ratio computation, subject-specific calibration, and threshold-based assignment of task-condition labels. The exact implementation details should be reported consistently with the firmware and analysis code.

### 3.1. EEG Hardware and Data Acquisition

**Fig. 2(a).**
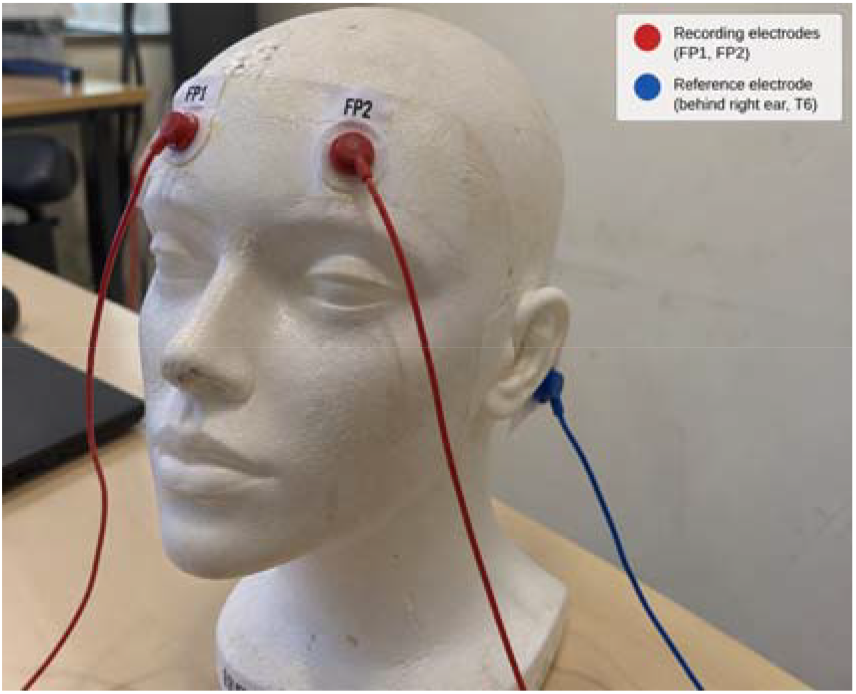
Electrode attachment during EEG acquisition.

**Fig. 2(b).**
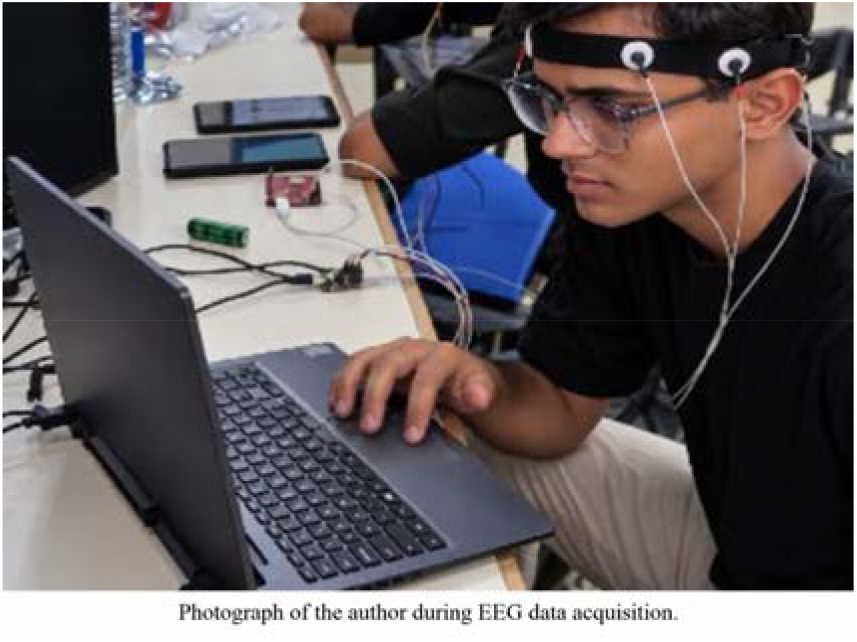
Experimental setup with acquisition hardware and computer.

**Fig. 2(c).**
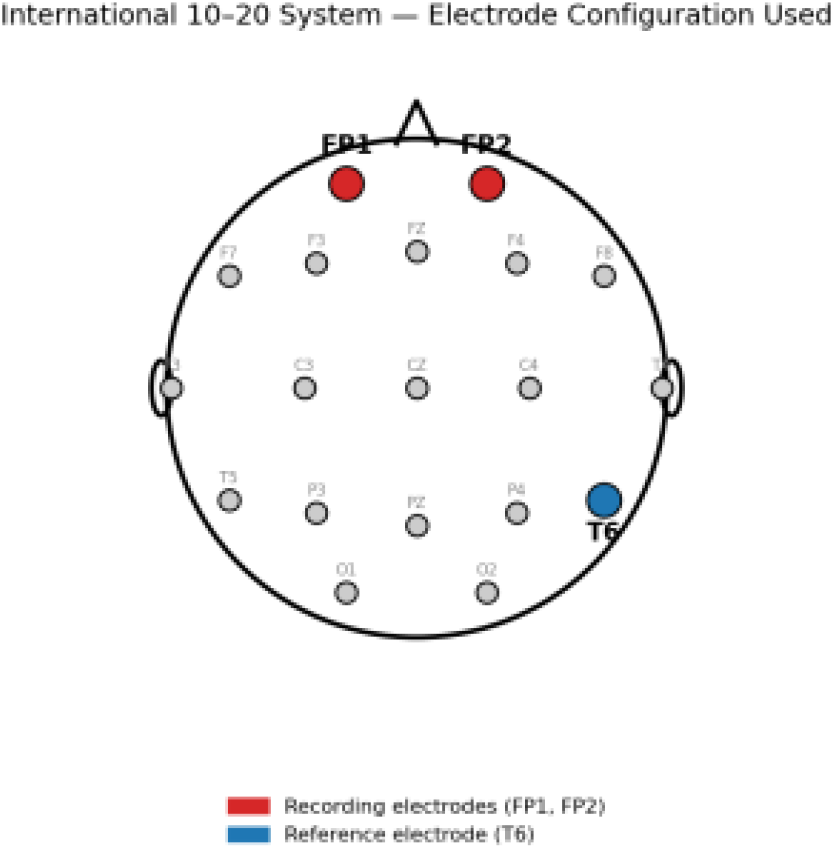
Electrode placement, including FP1, FP2 and the stated reference.

EEG data were acquired using a Brain BioAmp configured for two recording channels at FP1 and FP2. The stated sampling frequency was 256 Hz. The manuscript identifies T6 as the reference electrode; this placement and the reference/ground arrangement should be confirmed against the actual hardware wiring and acquisition configuration. The International 10–20 system was used to describe electrode placement.

Conductive electrode gel was applied and electrode contacts were secured. Recordings were made in an indoor laboratory, with participants seated and instructed to minimize movement, facial-muscle contractions, and excessive blinking. Three task conditions were recorded: Baseline (relaxed rest), Focused (sustained attention without time pressure), and Stressed (timed arithmetic under time pressure). The latter should be interpreted as a task condition intended to induce workload or stress, not as a validated psychological diagnosis.

For reproducibility, the final manuscript should report the number of participants, participant characteristics where appropriate, duration and number of trials per condition, exact ESP32 model, BioAmp version, electrode/reference/ground configuration, and any consent or ethics-review arrangements applicable to the study.

| Condition | Task description | Operational interpretation |
| --- | --- | --- |
| Baseline | Seated rest, normal breathing, no task. | Reference condition. |
| Focused | Sustained-attention task without time pressure. | Attention-demanding condition. |
| Stressed | Timed arithmetic under time pressure. | Workload/stress-induction condition; independent validation not reported. |

### 3.2. Signal Preprocessing

The manuscript reports removal of the DC offset, a 0.5–40 Hz band-pass filter, and a 50 Hz notch filter. The filter type, order, implementation location, and parameters should match the actual processing code and be stated explicitly. The recorded signals were visually inspected for movement artifacts and unstable electrode contact, and affected segments were excluded. If rejection was based on visual inspection, it should not be described as automated artifact rejection.

The signal was divided into 128-sample windows at 256 Hz. Each window therefore spans 0.5 s and has a nominal FFT-bin spacing of 2 Hz. The manuscript should specify whether a window function (for example, Hann) was applied before the FFT. If no window function was used, spectral leakage should be acknowledged.

### 3.3. Frequency-Domain Analysis Using the FFT

The Fast Fourier Transform (FFT) was used to obtain a frequency-domain representation of each analysis window. For a sampling frequency fs and a window containing N samples, the frequency associated with FFT bin k is:

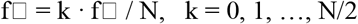

For fs = 256 Hz and N = 128, the nominal bin spacing is 2 Hz. The magnitude of the complex FFT coefficient is 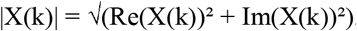. Magnitude is not itself power. If band power is calculated from squared FFT magnitudes, define the band estimate as P_B = Σ□□B |X(k)|^2^, with any required normalization stated. If the implementation instead uses a normalized power spectral density or another estimator, the equation and terminology must be changed to match that implementation. The final equations must be checked against the actual ESP32 and Python code before submission.

### 3.4. EEG Feature Extraction

The manuscript uses Theta/Alpha (θ/α) and Beta/Alpha (β/α) ratios as descriptive spectral features. The stated bands are Theta (4–8 Hz), Alpha (8–13 Hz), and Beta (13–30 Hz); the full plots also describe Delta and Gamma bands.

Band boundaries and FFT-bin assignment must be implemented consistently so that adjacent bands do not overlap unintentionally. The manuscript should identify the units and normalization of the band-power values, including whether they represent raw digital counts or calibrated physical units.

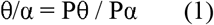

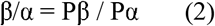

The ratios are relative spectral features and should not be interpreted as direct or uniquely specific measurements of psychological stress. The method should also specify how zero or very small Alpha power is handled to avoid unstable ratios.

### 3.5. Personalized Threshold-Based Classification

**Fig. 3.**
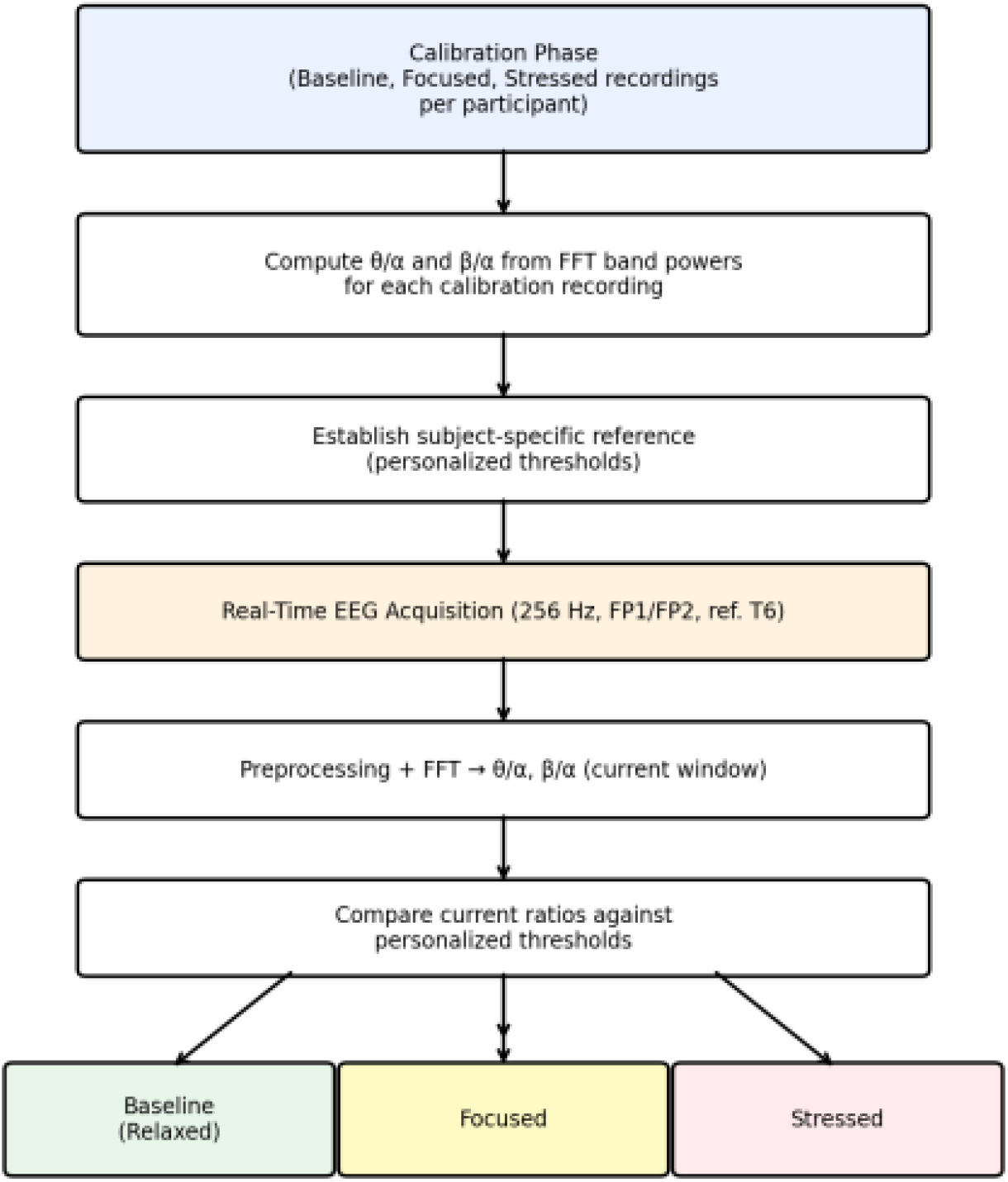
Personalized threshold-based classification workflow.

Let RB, RF, and RS denote the mean Beta/Alpha ratios calculated from the Baseline, Focused, and Stressed calibration recordings for a participant. The proposed midpoint thresholds are:

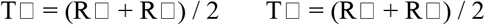

These thresholds assume that the calibration ratios are ordered and sufficiently separated. If this ordering is not observed, calibration should be treated as inconclusive rather than automatically assigning the intended labels. During monitoring, a Beta/Alpha value below T1 is assigned to Baseline, a value between T1 and T2 to Focused, and a value at or above T2 to Stressed only when the specified Theta/Alpha secondary criterion is also met. The final implementation must define the Theta/Alpha reference explicitly and document what happens when the secondary criterion is not met.

Because the thresholds are derived from calibration recordings, performance on those same recordings is not an independent estimate of classification accuracy. Independent recordings or held-out trials are needed to estimate generalization. The present method should therefore be described as a threshold-based feasibility framework until validated performance metrics are available.

## 4. Experimental Results and Discussion

The reported results compare spectral features across Baseline, Focused, and Stressed task conditions. Before final submission, all figures and table values should be regenerated or checked against the same verified analysis pipeline. The participant count, number of recordings, recording duration, and aggregation method used for each reported mean must be stated.

### 4.1. Time-Domain Band-Power Analysis

**Fig. 4 (a).**
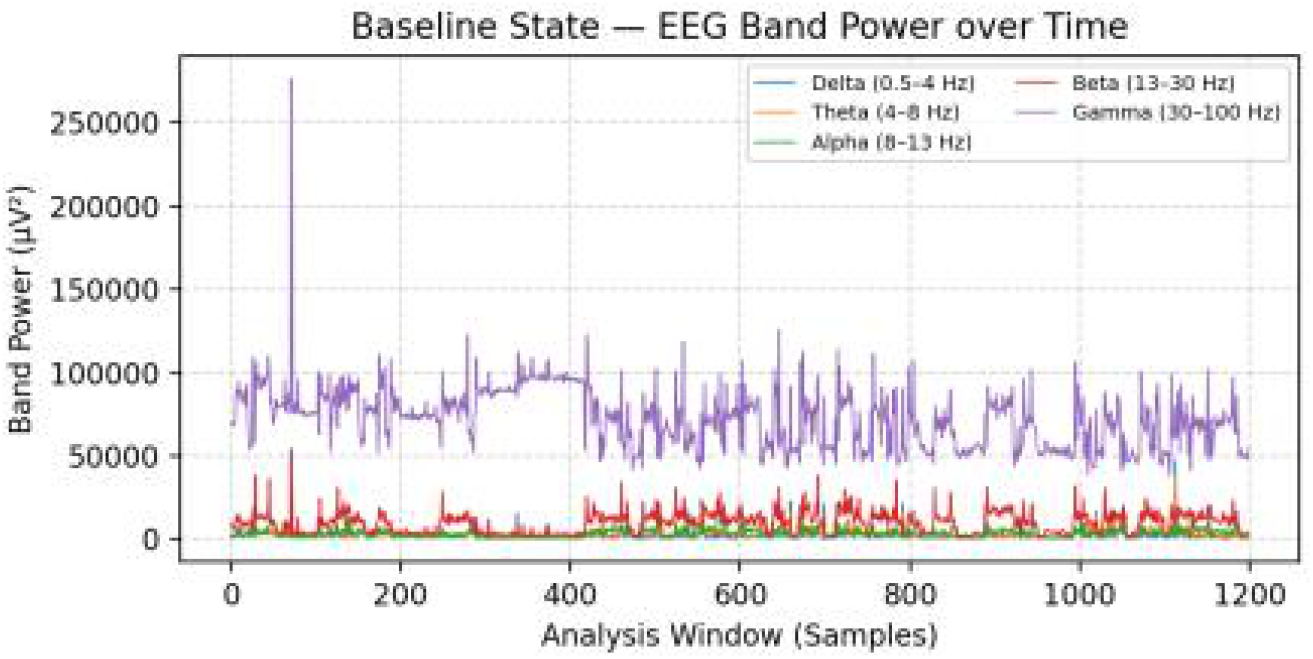
Baseline condition: EEG band power over analysis windows.

**Fig. 4 (b).**
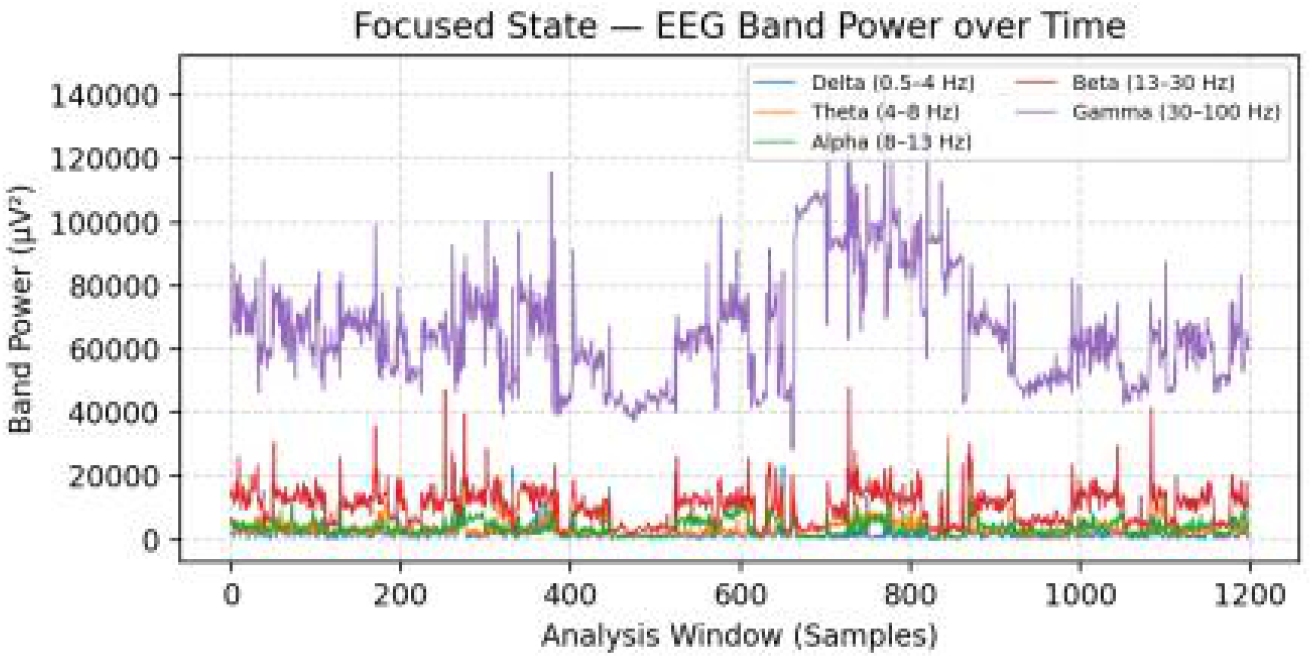
Focused condition: EEG band power over analysis windows.

**Fig. 4 (c).**
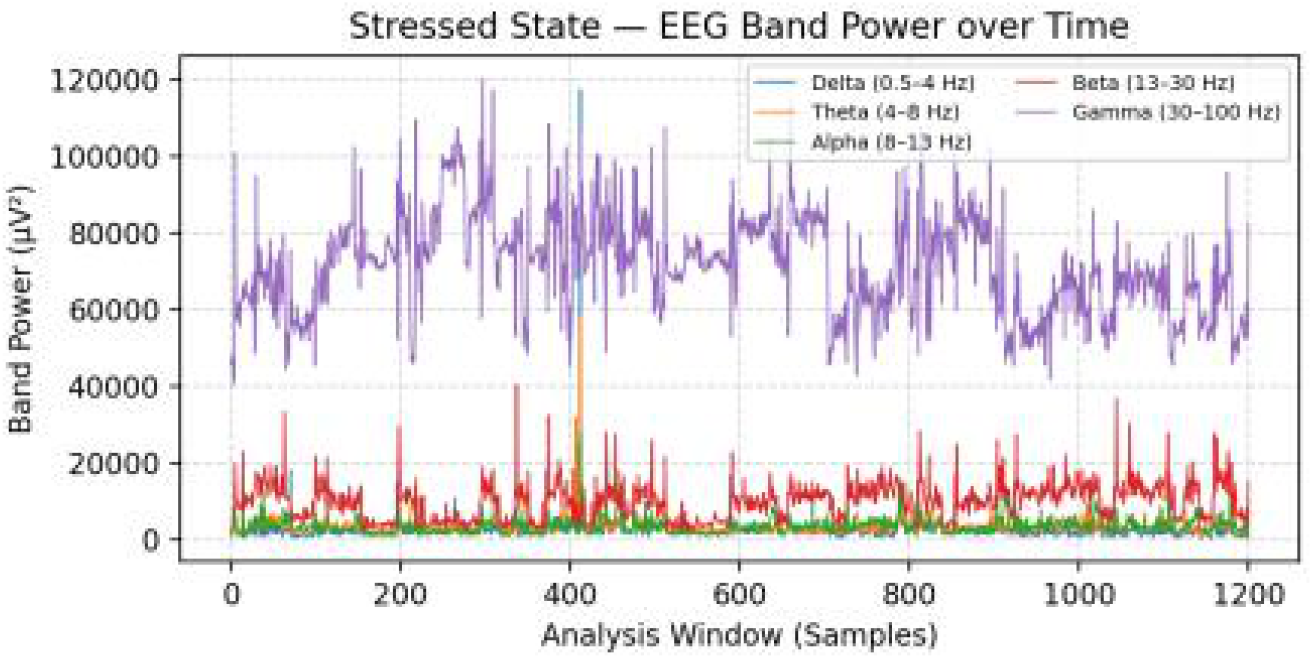
Stressed condition: EEG band power over analysis windows.

The reported mean Beta power is 9,976.14 during Baseline, 10,566.06 during Focused, and 9,885.59 during Stressed. Thus, Beta power is higher during Focused than Baseline, but lower during Stressed than during either of the other two conditions. The results do not support a monotonic increase in absolute Beta power across all three conditions. The figures should identify whether the horizontal axis represents time, sample number, or analysis-window index, and should include legible axis labels, units, and legends.

### 4.2. Frequency-Domain Analysis

**Fig. 5 (a).**
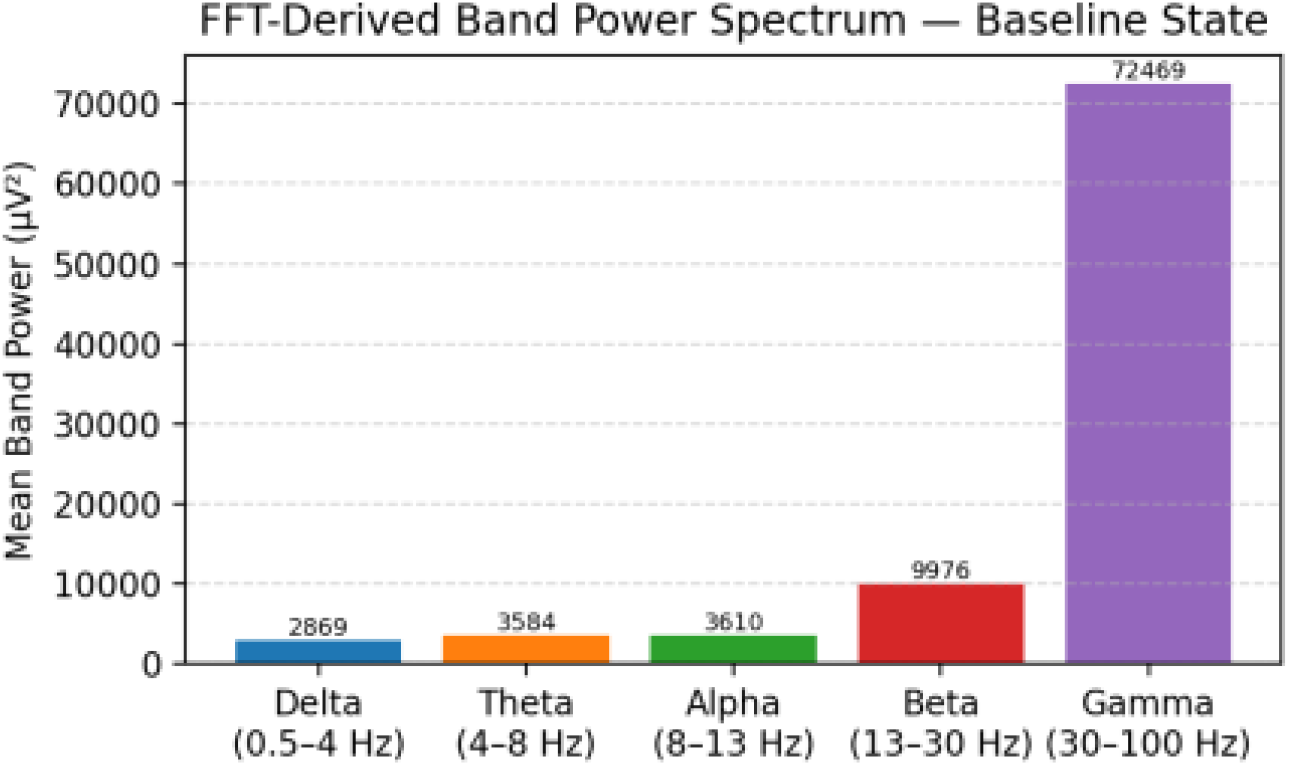
Baseline condition: FFT-derived band-power spectrum (original plotted data; verify Gamma range).

**Fig. 5 (b).**
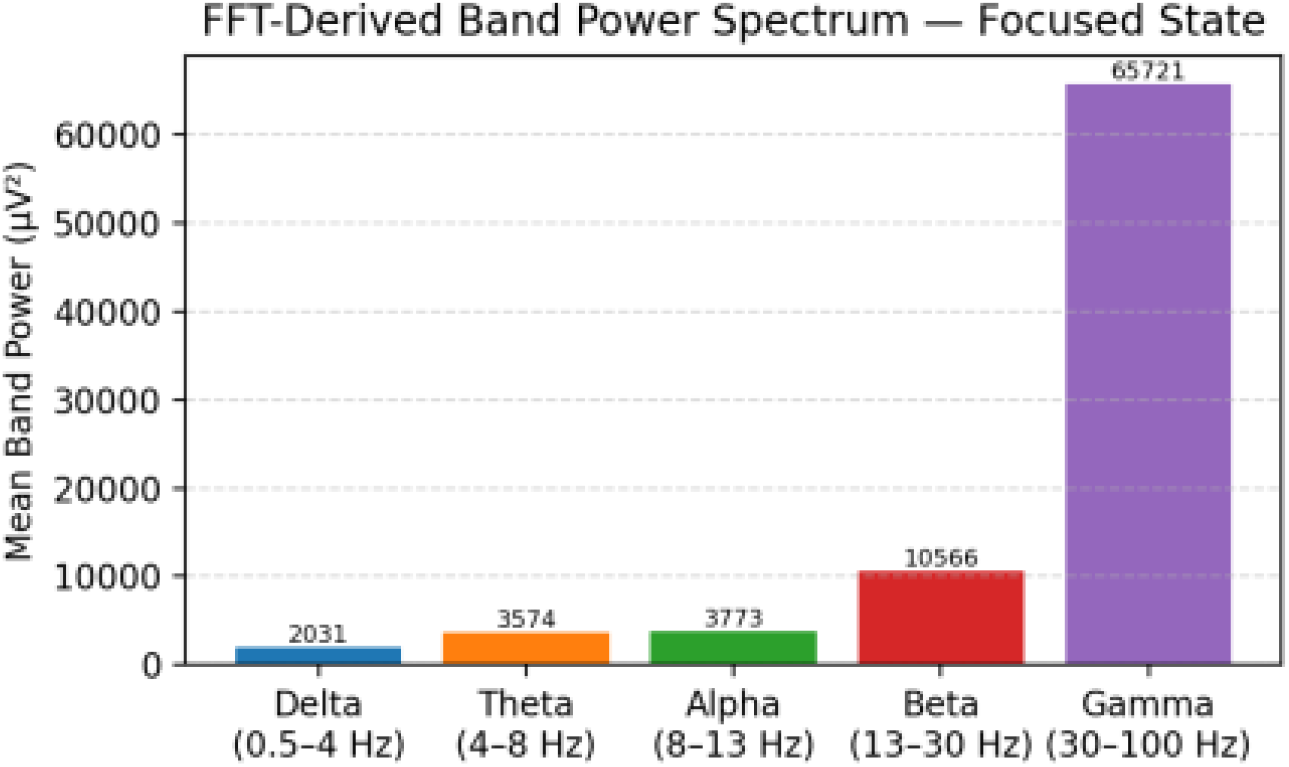
Focused condition: FFT-derived band-power spectrum (original plotted data; verify Gamma range).

**Fig. 5 (c).**
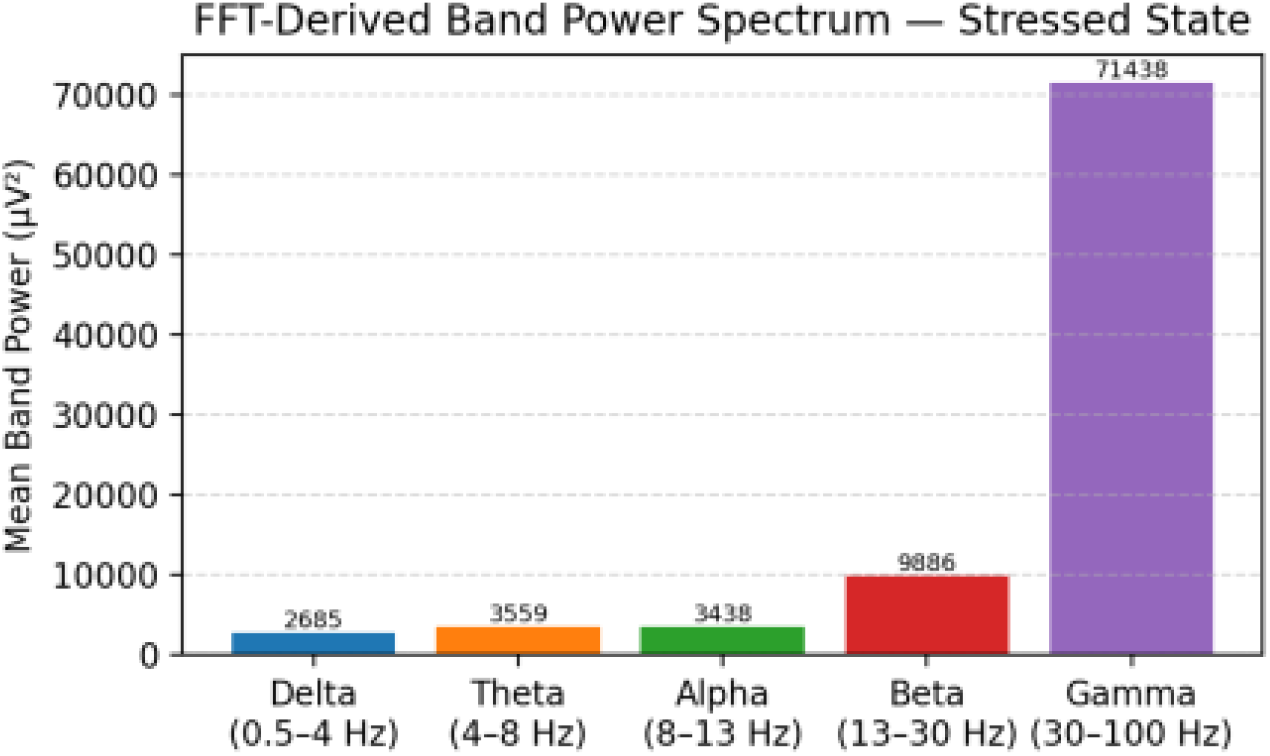
Stressed condition: FFT-derived band-power spectrum (original plotted data; verify Gamma range).

The stated preprocessing passband is 0.5–40 Hz. Accordingly, a Gamma band plotted from 30–100 Hz would be inconsistent with the described passband and must not be retained without evidence that the actual processing used a different passband. Verify the frequency bins and regenerate the figure from the underlying data. Confirm that all bands use a consistent power estimator and normalization, and explain whether plotted values are means across windows or values from a representative window.

The text should not claim that Beta power is the largest contributor across all plotted bands if the plotted Gamma values are larger. If the intended comparison concerns only Theta, Alpha, and Beta, state that scope explicitly.

### 4.3. Feature Extraction and Cognitive-State Analysis

The reported Beta/Alpha ratios are 2.76 for Baseline, 2.80 for Focused, and 2.88 for Stressed. The Baseline-to-Stressed difference is approximately 4.35%. The reported mean Alpha power decreases from 3,610.35 to 3,438.34, while mean Beta power also decreases slightly. Therefore, the higher Beta/Alpha ratio cannot be described as evidence of an absolute increase in Beta power.

**Fig. 6.**
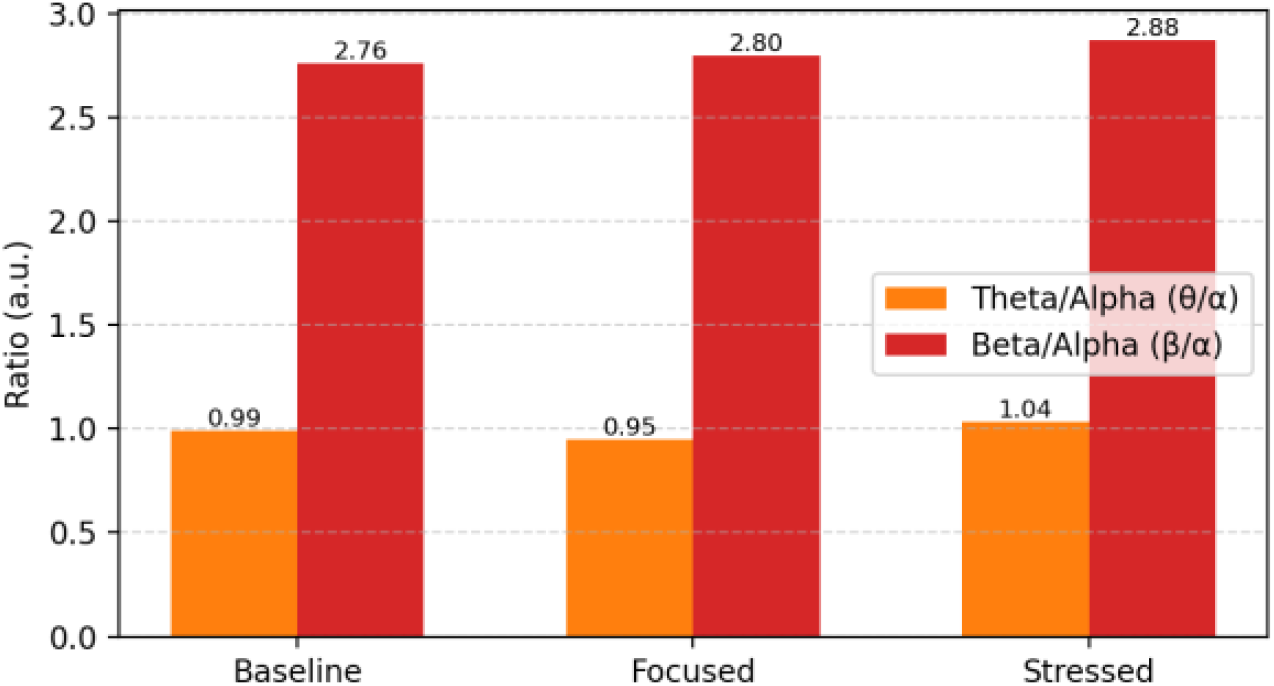
Comparison of EEG feature ratios across the three conditions (verify against final Table 2).

The reported Theta/Alpha ratios are 0.99 for Baseline, 0.95 for Focused, and 1.04 for Stressed. This pattern is not monotonic and should be reported descriptively. The manuscript must state whether each table ratio is calculated from the ratio of mean powers or from the mean of per-window ratios, because these calculations are not generally equivalent. Recalculate the table from the original data and ensure Figure 6 matches the final table.

| Condition | Mean Theta power | Mean Alpha power | Mean Beta power | θ/α ratio | β/α ratio |
| --- | --- | --- | --- | --- | --- |
| Baseline | 3584.15 | 3610.35 | 9976.14 | 0.99 | 2.76 |
| Focused | 3574.45 | 3772.69 | 10566.06 | 0.95 | 2.80 |
| Stressed | 3558.98 | 3438.34 | 9885.59 | 1.04 | 2.88 |

Note: The values above reproduce the reported table and have not been independently recalculated from raw recordings. Their units, aggregation method, and consistency with the original processing code must be confirmed before the table is treated as final.

### 4.4. Limitations

This is a feasibility pilot. The reported ratios are descriptive and are not accompanied by variance estimates or statistical tests. Classification accuracy, a confusion matrix, and a comparison with fixed thresholds are not reported. Timed arithmetic may reflect workload, time pressure, frustration, or other task effects, and an independent stress reference measure is not reported. FP1 and FP2 are susceptible to eye-blink and facial-muscle artifacts that may affect spectral estimates. Requiring a Stressed calibration recording may also be impractical in routine use. With 128 samples at 256 Hz, the nominal FFT-bin spacing is 2 Hz, limiting frequency resolution, particularly in low-frequency bands. Future evaluation should include additional participants, repeated sessions, documented artifact handling, independent validation, statistical analysis, and a biofeedback evaluation.

## 5. Conclusion

This study presents a low-cost, two-channel EEG processing framework that uses FFT-derived spectral features and subject-specific thresholds to distinguish Baseline, Focused, and Stressed task conditions. The reported pilot values show differences in the Beta/Alpha and Theta/Alpha ratios across the conditions. However, the magnitude and reliability of these differences require verification against the original processing pipeline and further evaluation.

Classification accuracy, statistical significance, and generalization across participants have not been established. Future work should include repeated recordings, independent validation, artifact handling, comparison with fixed thresholds, and evaluation of real-time biofeedback.

## Supporting information

App Used to take the Observation

## Data Availability

The original EEG recordings and derived band-power data are available from the authors upon request. Retain this statement only if the authors can provide the data and doing so is consistent with participant consent and applicable privacy requirements.

