## Supplementary material for "A Personalized Approach to Student Mental Health: Real-Time EEG Biofeedback for Stress and Focus Management": App Used to take the Observation: Main.py

```
import sys
import os
import time
import csv
import json
import pickle
import numpy as np
import pandas as pd
import serial
import serial.tools.list_ports
import random
from collections import deque
from scipy.fft import fft
from scipy.stats import entropy
from datetime import datetime

# GUI Imports
from PyQt6.QtWidgets import (QApplication, QMainWindow, QWidget, QVBoxLayout, QHBoxLayout, 
                             QPushButton, QLabel, QComboBox, QProgressBar, QStackedWidget, 
                             QTextEdit, QMessageBox, QFrame, QGridLayout, QScrollArea, 
                             QLineEdit, QFormLayout, QListWidget, QListWidgetItem, QSizePolicy, 
                             QSpacerItem, QButtonGroup, QRadioButton, QTabWidget, QGraphicsOpacityEffect, QDialog)
from PyQt6.QtCore import Qt, QThread, pyqtSignal, QTimer, QSize, QPropertyAnimation, QEasingCurve, QAbstractAnimation, QPoint
from PyQt6.QtGui import QFont, QIcon, QColor, QPalette, QCursor, QPainter, QPen, QBrush, QRadialGradient

# Plotting
import pyqtgraph as pg

# AI / Logic Imports
from sklearn.ensemble import RandomForestClassifier
from sklearn.preprocessing import LabelEncoder, StandardScaler
import joblib

# =================================================================
# GLOBAL CONFIGURATION
# =================================================================
SAMPLING_RATE = 256     # Hz
FFT_WINDOW = 64         # Samples
SEQUENCE_LENGTH = 10    # Temporal Window
STEP_SIZE = 32          # Overlap Step

# FILE PATHS
BASE_DIR = os.getcwd()

# Helper to get user-specific paths
def get_user_paths(username):
    user_dir = os.path.join(BASE_DIR, "Resource files", "With LSTM", "profiles", username)
    if not os.path.exists(user_dir):
        os.makedirs(user_dir)
    return {
        "RAW": os.path.join(user_dir, "Raw_EEG_Training_Data.csv"),
        "HISTORY": os.path.join(user_dir, "Session_History_Log.csv"),
        "MODEL": os.path.join(user_dir, "eeg_rf_model.pkl"),
        "SCALER": os.path.join(user_dir, "eeg_scaler.pkl"),
        "ENCODER": os.path.join(user_dir, "eeg_encoder.pkl"),
        "STATS": os.path.join(user_dir, "calibration_stats.pkl"),
        "PROFILE": os.path.join(user_dir, "user_profile.json")
    }

RAW_TRAINING_HEADERS = ["Label", "Raw_Value", "Timestamp"]

def extract_sequence_features(sequence):
    """Extract statistical features from a sequence for Random Forest.
    Converts sequence of band values into feature vector with mean, std, min, max, median."""
    features = []
    seq_array = np.array(sequence)
    
    for band_idx in range(seq_array.shape[1]):  # For each band
        band_values = seq_array[:, band_idx]
        features.extend([
            np.mean(band_values),      # Mean
            np.std(band_values),       # Standard Deviation
            np.min(band_values),       # Min
            np.max(band_values),       # Max
            np.median(band_values),    # Median
            np.max(band_values) - np.min(band_values)  # Range
        ])
    
    return features

def ensure_raw_training_headers(file_path):
    if not os.path.exists(file_path):
        with open(file_path, 'w', newline='') as f:
            csv.writer(f).writerow(RAW_TRAINING_HEADERS)
        return

    try:
        with open(file_path, 'r', newline='') as f:
            reader = csv.reader(f)
            first_row = next(reader, None)

            if first_row is None:
                with open(file_path, 'w', newline='') as fw:
                    csv.writer(fw).writerow(RAW_TRAINING_HEADERS)
                return

            if first_row == RAW_TRAINING_HEADERS:
                return

            header_set = set(first_row)
            backup = file_path + f".bak_{datetime.now().strftime('%Y%m%d_%H%M%S')}"
            temp_path = file_path + ".tmp"

            if header_set.issuperset(RAW_TRAINING_HEADERS):
                f.seek(0)
                dict_reader = csv.DictReader(f)
                rows = list(dict_reader)
                with open(temp_path, 'w', newline='') as fw:
                    writer = csv.writer(fw)
                    writer.writerow(RAW_TRAINING_HEADERS)
                    for row in rows:
                        writer.writerow([
                            row.get("Label", ""),
                            row.get("Raw_Value", ""),
                            row.get("Timestamp", "")
                        ])
                os.replace(file_path, backup)
                os.replace(temp_path, file_path)
                return

            if len(first_row) == 3 and not any(h in first_row for h in RAW_TRAINING_HEADERS):
                with open(temp_path, 'w', newline='') as fw:
                    writer = csv.writer(fw)
                    writer.writerow(RAW_TRAINING_HEADERS)
                    writer.writerow(first_row)
                    for row in reader:
                        writer.writerow(row)
                os.replace(file_path, backup)
                os.replace(temp_path, file_path)
                return

            os.replace(file_path, backup)
            with open(file_path, 'w', newline='') as fw:
                csv.writer(fw).writerow(RAW_TRAINING_HEADERS)
    except Exception:
        pass

CURRENT_USER_PATHS = {} # Will be set on login

# =================================================================
# LUXE BLACK & GOLD THEME
# =================================================================
THEME_GOLD = "#D4AF37"   
THEME_TEAL = "#00BCD4"   
THEME_RED  = "#FF5252"   
BG_DARK    = "#121212"   
PANEL_BG   = "#1E1E1E"   

STYLESHEET = f"""
QMainWindow {{ background-color: {BG_DARK}; }}
QWidget {{ font-family: 'Segoe UI', 'Inter', sans-serif; color: #E6E6E6; font-size: 13px; }}

QLabel#Title {{ font-size: 24px; font-weight: 700; color: {THEME_GOLD}; letter-spacing: 1px; }}
QLabel#Subtitle {{ font-size: 12px; color: #9AA0A6; letter-spacing: 1px; }}

/* Panels & Cards */
QFrame {{ 
    background-color: {PANEL_BG}; 
    border-radius: 12px; 
    border: 1px solid #2A2A2A;
}}
QFrame#NavPanel {{ 
    border-right: 1px solid #222; 
    background-color: #141418; 
    border-radius: 0px; 
}}
QFrame#Card {{ background: #141417; border: 1px solid #2C2C2C; }}
QFrame#Card:hover {{ border: 1px solid {THEME_GOLD}; }}

/* Tabs */
QTabWidget::pane {{ border: 1px solid #2A2A2A; background: {BG_DARK}; border-radius: 8px; }}
QTabBar::tab {{
    background: #1B1C21;
    color: #9AA0A6;
    padding: 10px 18px;
    border-top-left-radius: 8px;
    border-top-right-radius: 8px;
    margin-right: 4px;
}}
QTabBar::tab:selected {{ background: {THEME_GOLD}; color: #111; font-weight: 700; }}
QTabBar::tab:hover {{ background: #272830; color: #FFF; }}

/* Buttons */
QPushButton {{
    background-color: #1B1C21;
    border: 1px solid {THEME_GOLD};
    color: {THEME_GOLD};
    border-radius: 10px;
    padding: 10px 16px;
    font-weight: 700;
    letter-spacing: 0.8px;
}}
QPushButton:hover {{ background-color: {THEME_GOLD}; color: #121212; }}
QPushButton:pressed {{ background-color: #B08D55; border-color: #B08D55; }}
QPushButton:disabled {{ border: 1px solid #333; color: #555; background-color: #141414; }}

QPushButton#DangerBtn {{ border: 1px solid {THEME_RED}; color: {THEME_RED}; background-color: #1B1516; }}
QPushButton#DangerBtn:hover {{ background-color: {THEME_RED}; color: #FFF; }}

QPushButton#MenuBtn {{
    text-align: left; padding: 12px 14px; border: none; background: transparent; color: #A0A0A0; border-radius: 8px;
}}
QPushButton#MenuBtn:hover {{ color: #FFF; background: rgba(255,255,255,0.05); }}
QPushButton#MenuBtn:checked {{ 
    color: {THEME_GOLD}; 
    border-left: 4px solid {THEME_GOLD}; 
    background-color: rgba(212, 175, 55, 0.10); 
}}

/* Inputs */
QLineEdit, QTextEdit, QComboBox {{
    background-color: #0B0C10;
    border: 1px solid #30323A;
    color: #EEE;
    padding: 10px 12px;
    border-radius: 8px;
}}
QLineEdit:focus, QTextEdit:focus, QComboBox:focus {{ border: 1px solid {THEME_TEAL}; }}
QComboBox::drop-down {{ border: none; width: 24px; }}
QComboBox QAbstractItemView {{ background: #15161B; border: 1px solid #2C2C2C; selection-background-color: #2E2E2E; }}

/* Progress Bar */
QProgressBar {{ border: none; background-color: #2A2A2A; border-radius: 6px; height: 12px; text-align: center; }}
QProgressBar::chunk {{ background-color: {THEME_TEAL}; border-radius: 6px; }}

/* Scroll Area */
QScrollArea {{ background: transparent; border: none; }}
QScrollBar:vertical {{ background: transparent; width: 10px; margin: 4px; }}
QScrollBar::handle:vertical {{ background: #2E2F36; border-radius: 5px; min-height: 24px; }}
QScrollBar::handle:vertical:hover {{ background: #3A3C44; }}
QScrollBar::add-line:vertical, QScrollBar::sub-line:vertical {{ height: 0px; }}
QScrollBar:horizontal {{ background: transparent; height: 10px; margin: 4px; }}
QScrollBar::handle:horizontal {{ background: #2E2F36; border-radius: 5px; min-width: 24px; }}
QScrollBar::handle:horizontal:hover {{ background: #3A3C44; }}
QScrollBar::add-line:horizontal, QScrollBar::sub-line:horizontal {{ width: 0px; }}

/* Lists */
QListWidget {{ background: #0B0C10; border: 1px solid #2C2C2C; border-radius: 8px; padding: 8px; }}
QListWidget::item {{ padding: 6px 4px; }}
QListWidget::item:selected {{ background: #2B2C33; color: #FFF; border-radius: 6px; }}
"""

# =================================================================
# ADVANCED SIGNAL PROCESSING LOGIC
# =================================================================

def compute_features(raw_window):
    """
    Advanced Feature Extraction: 
    1. FFT (5 Bands)
    2. Spectral Entropy (Complexity)
    3. Hjorth Activity (Variance)
    Returns vector of size 7.
    """
    try:
        # 1. Artifact Removal Check (Simple Amplitude Threshold)
        if np.max(raw_window) > 4095 or np.min(raw_window) < 0:
            return [0] * 7

        N = len(raw_window)
        # Apply Hanning window
        windowed_sig = raw_window * np.hanning(N)
        yf = fft(windowed_sig)
        xf = np.linspace(0.0, SAMPLING_RATE/2, N//2)
        mags = 2.0/N * np.abs(yf[0:N//2])
        
        # Normalize mags for entropy
        psd_norm = mags / (np.sum(mags) + 1e-9)
        
        # --- Standard Bands ---
        d = np.mean(mags[(xf >= 0.5) & (xf < 4)])
        t = np.mean(mags[(xf >= 4) & (xf < 8)])
        a = np.mean(mags[(xf >= 8) & (xf < 13)])
        b = np.mean(mags[(xf >= 13) & (xf < 30)])
        g = np.mean(mags[(xf >= 30) & (xf < 45)])
        
        # --- Advanced Features ---
        spec_entropy = entropy(psd_norm)
        if np.isnan(spec_entropy): spec_entropy = 0
        
        activity = np.var(raw_window)
        
        return [d, t, a, b, g, spec_entropy, activity]
        
    except Exception:
        return [0, 0, 0, 0, 0, 0, 0]

def generate_demo_dataset():
    """Generates synthetic EEG data with ORGANIC VARIANCE for better RF training."""
    print("Generating Synthetic Data...")
    data_rows = []
    start_time = datetime.now()
    duration = 120 # 2 mins per state for demo
    t = np.linspace(0, duration, int(duration * SAMPLING_RATE), endpoint=False)
    dc = 2048 
    
    # 1. Baseline: Moderate Alpha, low Beta. Steady "breathing" rhythm.
    am_base = 1.0 + 0.3 * np.sin(2 * np.pi * 0.2 * t) 
    noise = np.random.normal(0, 15, len(t))
    sig = (100 * np.sin(2*np.pi*10*t) + 30 * np.sin(2*np.pi*20*t)) * am_base
    base_sig = np.clip(dc + sig + noise, 0, 4095).astype(int)
    
    # 2. Stressed: High Beta, Spiky. Fast, erratic modulation.
    am_stress = 1.0 + 0.5 * np.random.rand(len(t)) # Jittery amplitude
    noise = np.random.normal(0, 40, len(t)) # More noise
    sig = (30 * np.sin(2*np.pi*10*t) + 300 * np.sin(2*np.pi*25*t)) * am_stress
    stress_sig = np.clip(dc + sig + noise, 0, 4095).astype(int)
    
    # 3. Focused: Very Strong Alpha, Low Theta. Steady flow.
    am_focus = 1.0 # Steady focus
    noise = np.random.normal(0, 10, len(t)) # Low noise
    sig = (350 * np.sin(2*np.pi*10*t) + 20 * np.sin(2*np.pi*5*t)) * am_focus
    focus_sig = np.clip(dc + sig + noise, 0, 4095).astype(int)
    
    for val in base_sig: data_rows.append(["Baseline", val, start_time.strftime("%Y-%m-%d %H:%M:%S")])
    for val in stress_sig: data_rows.append(["Stressed", val, start_time.strftime("%Y-%m-%d %H:%M:%S")])
    for val in focus_sig: data_rows.append(["Focused", val, start_time.strftime("%Y-%m-%d %H:%M:%S")])
            
    df = pd.DataFrame(data_rows, columns=["Label", "Raw_Value", "Timestamp"])
    df.to_csv(CURRENT_USER_PATHS["RAW"], index=False)
    return len(df)

# =================================================================
# BACKGROUND THREADS
# =================================================================

class RecorderThread(QThread):
    progress_update = pyqtSignal(int, int, int) 
    live_signal = pyqtSignal(float) 
    feedback_msg = pyqtSignal(str, str) 
    finished_phase = pyqtSignal(str)
    error_occurred = pyqtSignal(str)
    signal_quality = pyqtSignal(bool)
    connection_status = pyqtSignal(str, str)
    signal_strength = pyqtSignal(int)

    def __init__(self, port, label, duration=1200):
        super().__init__()
        self.port = port
        self.label = label
        self.duration = duration
        self.running = True
        self.signal_buffer = deque(maxlen=256) 
        self.last_data_time = time.time()
        self.no_data_emitted = False

    def run(self):
        ser = None
        f = None
        try:
            ser = serial.Serial(self.port, 115200, timeout=1)
            time.sleep(2); ser.flushInput()
            self.connection_status.emit("CONNECTED", THEME_TEAL)
            
            start_time = time.time()
            count = 0
            
            f_path = CURRENT_USER_PATHS["RAW"]
            ensure_raw_training_headers(f_path)
            mode = 'a' if os.path.exists(f_path) else 'w'
            f = open(f_path, mode, newline='')
            writer = csv.writer(f)
            
            if mode == 'w': writer.writerow(RAW_TRAINING_HEADERS)
            
            while self.running:
                elapsed = time.time() - start_time
                if elapsed >= self.duration: break
                
                if ser.in_waiting:
                    try:
                        line = ser.readline().decode('utf-8', errors='ignore').strip()
                        if line.isdigit():
                            val = int(line)
                            self.last_data_time = time.time()
                            if self.no_data_emitted:
                                self.connection_status.emit("CONNECTED", THEME_TEAL)
                                self.no_data_emitted = False
                            
                            self.signal_buffer.append(val)
                            self.live_signal.emit(val)
                            if count % 10 == 0:
                                self.signal_quality.emit(10 < val < 4090)
                            
                            if count % 64 == 0: 
                                self.check_quality()
                            
                            timestamp = datetime.now().strftime("%Y-%m-%d %H:%M:%S")
                            writer.writerow([self.label, line, timestamp])
                            
                            if count % 256 == 0: f.flush()
                            count += 1
                            
                            if count % 128 == 0:
                                pct = int((elapsed / self.duration) * 100)
                                rem = int(self.duration - elapsed)
                                self.progress_update.emit(pct, rem, count)
                    except: pass
                else:
                    if time.time() - self.last_data_time > 3 and not self.no_data_emitted:
                        self.feedback_msg.emit("⚠️ NO DATA STREAM. Check cable / sensor.", THEME_RED)
                        self.connection_status.emit("NO DATA", THEME_RED)
                        self.signal_quality.emit(False)
                        self.signal_strength.emit(0)
                        self.no_data_emitted = True
                    time.sleep(0.01)
            
            self.finished_phase.emit(self.label)
            
        except Exception as e:
            self.connection_status.emit("DISCONNECTED", THEME_RED)
            self.error_occurred.emit(f"Connection Error: {str(e)}")
        finally:
            if ser and ser.is_open: ser.close()
            if f: f.close()
            self.connection_status.emit("DISCONNECTED", "#777")

    def check_quality(self):
        if len(self.signal_buffer) < 50: return
        data = np.array(self.signal_buffer)
        std = np.std(data)
        mn, mx = np.min(data), np.max(data)
        
        if mx > 4090 or mn < 10:
            self.feedback_msg.emit("⚠️ SIGNAL CLIPPING! Check Connections.", THEME_RED)
            self.signal_quality.emit(False)
            self.signal_strength.emit(5)
        elif std < 5:
            self.feedback_msg.emit("⚠️ SIGNAL FLATLINE. Is device on?", THEME_RED)
            self.signal_quality.emit(False)
            self.signal_strength.emit(10)
        elif std > 500:
            self.feedback_msg.emit("⚠️ HIGH NOISE (EMG). Relax muscles.", "#FF9800")
            self.signal_quality.emit(False)
            self.signal_strength.emit(35)
        else:
            self.feedback_msg.emit("✅ SIGNAL GOOD. Keep steady.", "#00FF00")
            self.signal_quality.emit(True)
            std_norm = np.clip((std - 10) / (200 - 10), 0, 1)
            score = int(60 + std_norm * 40)
            self.signal_strength.emit(score)

    def stop(self):
        self.running = False

class TrainerThread(QThread):
    log_update = pyqtSignal(str)
    training_finished = pyqtSignal()

    def run(self):
        self.log_update.emit(">>> INITIATING TRAINING PROTOCOL...")
        f_path = CURRENT_USER_PATHS["RAW"]
        if not os.path.exists(f_path):
            self.log_update.emit("ERROR: No Neural Data Found.")
            self.training_finished.emit()
            return

        ensure_raw_training_headers(f_path)

        try:
            try:
                df = pd.read_csv(f_path, on_bad_lines='skip')
            except:
                self.log_update.emit("CRITICAL ERROR: Data Corrupted.")
                self.training_finished.emit()
                return

            features, labels = [], []
            raw_bands = {'Baseline': [], 'Stressed': [], 'Focused': []}

            self.log_update.emit(">>> EXTRACTING FEATURES...")
            
            for label, group in df.groupby('Label'):
                raw = pd.to_numeric(group['Raw_Value'], errors='coerce').dropna().values
                if len(raw) < FFT_WINDOW: continue
                
                label_feats = []
                for i in range(0, len(raw) - FFT_WINDOW, STEP_SIZE):
                    win = raw[i:i+FFT_WINDOW]
                    bands = compute_features(win) 
                    label_feats.append(bands)
                
                # --- SMART OUTLIER REJECTION ---
                if len(label_feats) > 10:
                    data_np = np.array(label_feats)
                    mean = np.mean(data_np, axis=0)
                    std = np.std(data_np, axis=0)
                    std = np.where(std < 1e-9, 1.0, std) 
                    
                    clean_feats = []
                    for f in label_feats:
                        z_score = np.abs((f - mean) / std)
                        if np.all(z_score < 4): clean_feats.append(f)
                    
                    if len(clean_feats) < len(label_feats) * 0.1:
                        self.log_update.emit(f"  > {label}: Outlier logic too aggressive. Using raw data.")
                        clean_feats = label_feats
                        
                    for f in clean_feats:
                        features.append(f)
                        labels.append(label)
                        if label in raw_bands: raw_bands[label].append(f)
                    self.log_update.emit(f"  > {label}: Kept {len(clean_feats)}/{len(label_feats)} samples")
                else:
                    for f in label_feats:
                        features.append(f); labels.append(label)
            
            if not features:
                self.log_update.emit("ERROR: Insufficient Data.")
                self.training_finished.emit()
                return

            self.log_update.emit(f"DATASET: {len(features)} Tensors.")

            self.log_update.emit(">>> COMPUTING HEURISTICS...")
            if raw_bands['Baseline']:
                base = np.array(raw_bands['Baseline'])
                avg_a = np.mean(base[:, 2]); avg_b = np.mean(base[:, 3]); avg_t = np.mean(base[:, 1]) 
                stats = {'stress_ratio': avg_b/(avg_a+1e-9), 'focus_ratio': avg_a/(avg_t+1e-9), 'baseline_alpha': avg_a, 'baseline_beta': avg_b, 'baseline_theta': avg_t}
                with open(CURRENT_USER_PATHS["STATS"], 'wb') as f: pickle.dump(stats, f)
                self.log_update.emit(f"  [MATH] Baseline Calibrated.")
            
            self.log_update.emit(">>> TRAINING RANDOM FOREST...")
            encoder = LabelEncoder()
            y_enc = encoder.fit_transform(labels)
            
            scaler = StandardScaler()
            X_scaled = scaler.fit_transform(features)
            
            # Extract sequence-based features for RF
            X_seq_features = []
            y_seq = []
            for i in range(len(X_scaled) - SEQUENCE_LENGTH):
                seq_feature = extract_sequence_features(X_scaled[i:i+SEQUENCE_LENGTH])
                X_seq_features.append(seq_feature)
                y_seq.append(y_enc[i+SEQUENCE_LENGTH])
            
            X_seq_features = np.array(X_seq_features)
            y_seq = np.array(y_seq)
            
            # Train Random Forest with optimized parameters
            model = RandomForestClassifier(
                n_estimators=200,
                max_depth=20,
                min_samples_split=5,
                min_samples_leaf=2,
                max_features='sqrt',
                class_weight='balanced',
                random_state=42,
                n_jobs=-1
            )
            model.fit(X_seq_features, y_seq)
            
            # Calculate accuracy
            acc = model.score(X_seq_features, y_seq)
            
            joblib.dump(model, CURRENT_USER_PATHS["MODEL"])
            with open(CURRENT_USER_PATHS["SCALER"], 'wb') as f: pickle.dump(scaler, f)
            with open(CURRENT_USER_PATHS["ENCODER"], 'wb') as f: pickle.dump(encoder, f)
            
            self.log_update.emit(f">>> RANDOM FOREST CONVERGED. Accuracy: {acc*100:.1f}%")
            
        except Exception as e:
            self.log_update.emit(f"SYSTEM FAILURE: {str(e)}")
        finally:
            self.training_finished.emit()

class MonitorThread(QThread):
    data_update = pyqtSignal(list, str, float, str, float, float) 
    dashboard_update = pyqtSignal(str)
    signal_quality = pyqtSignal(bool)
    error_occurred = pyqtSignal(str)

    def __init__(self, port):
        super().__init__()
        self.port = port
        self.running = True
        self.last_logged_state = "None"
        self.last_log_time = time.time()
        self.signal_buffer = deque(maxlen=256)
        self.last_data_time = time.time()
        self.no_data_emitted = False

    def run(self):
        ser = None
        log_file = None
        try:
            if not os.path.exists(CURRENT_USER_PATHS["MODEL"]):
                self.error_occurred.emit("Model not found! Train AI first.")
                return

            model = joblib.load(CURRENT_USER_PATHS["MODEL"])
            with open(CURRENT_USER_PATHS["SCALER"], 'rb') as f: scaler = pickle.load(f)
            with open(CURRENT_USER_PATHS["ENCODER"], 'rb') as f: encoder = pickle.load(f)
            
            base_s = 1.0; base_f = 1.0; base_beta = 0.0; base_alpha = 0.0
            if os.path.exists(CURRENT_USER_PATHS["STATS"]):
                with open(CURRENT_USER_PATHS["STATS"], 'rb') as f: stats = pickle.load(f)
                base_s = stats.get('stress_ratio', 1.0)
                base_f = stats.get('focus_ratio', 1.0)
                base_beta = stats.get('baseline_beta', 0.0)
                base_alpha = stats.get('baseline_alpha', 0.0)
            
            ser = serial.Serial(self.port, 115200, timeout=1)
            time.sleep(2); ser.flushInput()
            
            hist_path = CURRENT_USER_PATHS["HISTORY"]
            mode = 'a' if os.path.exists(hist_path) else 'w'
            log_file = open(hist_path, mode, newline='')
            writer = csv.writer(log_file)
            if mode == 'w': writer.writerow(["Timestamp", "Event", "Confidence"])

            raw_buf = deque(maxlen=FFT_WINDOW)
            seq_buf = deque(maxlen=SEQUENCE_LENGTH)
            for _ in range(SEQUENCE_LENGTH): seq_buf.append([0.0]*7) 
            
            ui_throttle = 0 

            while self.running:
                if ser.in_waiting:
                    line = ser.readline().decode('utf-8', errors='ignore').strip()
                    if line.isdigit():
                        val = float(line)
                        raw_buf.append(val)
                        self.signal_buffer.append(val)
                        self.last_data_time = time.time()
                        if self.no_data_emitted:
                            self.signal_quality.emit(True)
                            self.no_data_emitted = False
                        
                        if ui_throttle % 64 == 0:
                            self.check_quality()

                        if len(raw_buf) == FFT_WINDOW:
                            feats = compute_features(list(raw_buf))
                            scaled = scaler.transform(np.array(feats).reshape(1,-1))[0]
                            seq_buf.append(scaled)
                            
                            # Extract sequence features for RF
                            seq_features = extract_sequence_features(list(seq_buf))
                            pred = model.predict(np.array(seq_features).reshape(1, -1))[0]
                            pred_proba = model.predict_proba(np.array(seq_features).reshape(1, -1))[0]
                            
                            label = encoder.inverse_transform([pred])[0]
                            conf = np.max(pred_proba)
                            
                            curr_s = feats[3]/(feats[2]+1e-9) 
                            curr_f = feats[2]/(feats[1]+1e-9) 
                            
                            tag = "MATCH"
                            final_state = "Neutral"
                            
                            if label == "Stressed":
                                if (curr_s > base_s * 1.1) or (feats[3] > base_beta * 1.2):
                                    tag = "VERIFIED"; final_state = "STRESSED"
                                else: tag = "UNVERIFIED"
                            
                            elif label == "Focused":
                                if (curr_f > base_f * 1.1) or (feats[2] > base_alpha * 1.2):
                                    tag = "VERIFIED"; final_state = "FOCUSED"
                                else: tag = "UNVERIFIED"
                            
                            elif label == "Baseline":
                                final_state = "CALM"
                                
                            self.data_update.emit(feats[:5], label, conf, tag, curr_s, curr_f) 
                            self.dashboard_update.emit(final_state)
                            
                            if final_state != "Neutral" and tag == "VERIFIED":
                                if final_state != self.last_logged_state or (time.time() - self.last_log_time > 5):
                                    t_str = datetime.now().strftime("%H:%M:%S")
                                    writer.writerow([t_str, f"State: {final_state}", f"{conf:.2f}"])
                                    log_file.flush()
                                    self.last_logged_state = final_state
                                    self.last_log_time = time.time()
                            
                            for _ in range(STEP_SIZE): raw_buf.popleft()
                            ui_throttle += 1
                else:
                    if time.time() - self.last_data_time > 3 and not self.no_data_emitted:
                        self.signal_quality.emit(False)
                        self.no_data_emitted = True
                    time.sleep(0.01)
                            
        except Exception as e:
            self.error_occurred.emit(str(e))
        finally:
            if ser and ser.is_open: ser.close()
            if log_file: log_file.close()

    def stop(self):
        self.running = False

    def check_quality(self):
        if len(self.signal_buffer) < 50:
            return
        data = np.array(self.signal_buffer)
        std = np.std(data)
        mn, mx = np.min(data), np.max(data)
        
        if mx > 4090 or mn < 10:
            self.signal_quality.emit(False)
        elif std < 5:
            self.signal_quality.emit(False)
        elif std > 500:
            self.signal_quality.emit(False)
        else:
            self.signal_quality.emit(True)

# =================================================================
# TASK WIDGETS
# =================================================================

class BreathingTask(QWidget):
    def __init__(self):
        super().__init__()
        layout = QVBoxLayout()
        self.lbl = QLabel("Breathe In")
        self.lbl.setAlignment(Qt.AlignmentFlag.AlignCenter)
        self.lbl.setStyleSheet(f"font-size: 40px; font-weight: bold; color: {THEME_TEAL};")
        layout.addWidget(self.lbl)
        
        opt_lay = QHBoxLayout()
        self.btn_478 = QRadioButton("4-7-8 (Calm)"); self.btn_478.setChecked(True)
        self.btn_box = QRadioButton("Box (Focus)")
        
        self.btn_478.toggled.connect(self.update_mode)
        self.btn_box.toggled.connect(self.update_mode)
        
        opt_lay.addStretch(); opt_lay.addWidget(self.btn_478); opt_lay.addWidget(self.btn_box); opt_lay.addStretch()
        layout.addLayout(opt_lay)
        
        self.setLayout(layout)
        self.timer = QTimer(); self.timer.timeout.connect(self.tick); self.step = 0; self.mode = "4-7-8"

    def start(self): 
        self.update_mode()
        self.step = 0; self.timer.start(1000)
    
    def update_mode(self):
        self.mode = "box" if self.btn_box.isChecked() else "4-7-8"
        
    def stop(self): self.timer.stop(); self.lbl.setText("Relax")
    def get_score(self): return self.step * 10
    
    def tick(self):
        self.step += 1
        cycle = 16 if self.mode == "box" else 19
        curr = self.step % cycle
        if self.mode == "box":
            if curr < 4: self.lbl.setText(f"INHALE ({4-curr})")
            elif curr < 8: self.lbl.setText(f"HOLD ({8-curr})")
            elif curr < 12: self.lbl.setText(f"EXHALE ({12-curr})")
            else: self.lbl.setText(f"HOLD ({16-curr})")
        else:
            if curr < 4: self.lbl.setText(f"INHALE ({4-curr})")
            elif curr < 11: self.lbl.setText(f"HOLD ({11-curr})")
            else: self.lbl.setText(f"EXHALE ({19-curr})")

class StroopTask(QWidget):
    def __init__(self):
        super().__init__()
        layout = QVBoxLayout()
        self.word_lbl = QLabel("BLUE")
        self.word_lbl.setAlignment(Qt.AlignmentFlag.AlignCenter)
        self.word_lbl.setStyleSheet("font-size: 60px; font-weight: bold; margin: 20px;")
        layout.addWidget(self.word_lbl)
        
        mode_lay = QHBoxLayout()
        self.mode_stroop = QRadioButton("Stroop"); self.mode_stroop.setChecked(True)
        self.mode_math = QRadioButton("Rapid Math")
        self.mode_stroop.toggled.connect(self.switch_mode)
        self.mode_math.toggled.connect(self.switch_mode)
        mode_lay.addStretch(); mode_lay.addWidget(self.mode_stroop); mode_lay.addWidget(self.mode_math); mode_lay.addStretch()
        layout.addLayout(mode_lay)
        
        self.stroop_widget = QWidget()
        btn_layout = QHBoxLayout()
        self.colors = ["RED", "BLUE", "GREEN", "YELLOW"]
        self.color_map = {"RED": "#FF0000", "BLUE": "#0000FF", "GREEN": "#00FF00", "YELLOW": "#FFFF00"}
        for c in self.colors:
            btn = QPushButton(c); btn.setStyleSheet(f"background: {self.color_map[c]}; color: black; font-weight: bold; padding: 20px; border: none;")
            btn.clicked.connect(lambda _, x=c: self.check_stroop(x))
            btn_layout.addWidget(btn)
        self.stroop_widget.setLayout(btn_layout)
        layout.addWidget(self.stroop_widget)
        
        self.math_widget = QWidget(); self.math_widget.hide()
        ml = QVBoxLayout()
        self.math_in = QLineEdit(); self.math_in.setPlaceholderText("Answer")
        self.math_in.returnPressed.connect(self.check_math)
        ml.addWidget(self.math_in)
        self.math_widget.setLayout(ml)
        layout.addWidget(self.math_widget)
        
        self.score_lbl = QLabel("Score: 0"); self.score_lbl.setAlignment(Qt.AlignmentFlag.AlignCenter)
        layout.addWidget(self.score_lbl)
        self.setLayout(layout)
        self.score = 0; self.timer = QTimer(); self.timer.timeout.connect(self.next_round)
        self.is_math = False

    def switch_mode(self):
        self.is_math = self.mode_math.isChecked()
        if self.is_math:
            self.stroop_widget.hide(); self.math_widget.show()
        else:
            self.stroop_widget.show(); self.math_widget.hide()
        self.next_round()

    def start(self): 
        self.score = 0; self.score_lbl.setText("Score: 0")
        self.switch_mode() 
        self.timer.start(3000) 

    def stop(self): self.timer.stop()
    def get_score(self): return self.score

    def next_round(self):
        if self.is_math:
            a, b = random.randint(10, 99), random.randint(10, 99)
            op = random.choice(["+", "-"])
            self.math_ans = a + b if op == "+" else a - b
            self.word_lbl.setText(f"{a} {op} {b} = ?")
            self.word_lbl.setStyleSheet(f"font-size: 60px; font-weight: bold; color: {THEME_RED};")
            self.math_in.clear(); self.math_in.setFocus()
        else:
            word = random.choice(self.colors); ink = random.choice(self.colors); self.current_ink = ink
            self.word_lbl.setText(word)
            self.word_lbl.setStyleSheet(f"font-size: 60px; font-weight: bold; color: {self.color_map[ink]}; margin: 20px;")

    def check_stroop(self, color_name):
        if color_name == self.current_ink: self.score += 50; self.score_lbl.setStyleSheet(f"color: {THEME_TEAL}; font-size: 18px;")
        else: self.score_lbl.setStyleSheet("color: red; font-size: 18px;")
        self.score_lbl.setText(f"Score: {self.score}"); self.next_round(); self.timer.start(3000)

    def check_math(self):
        try:
            if int(self.math_in.text()) == self.math_ans: self.score += 50; self.score_lbl.setStyleSheet(f"color: {THEME_TEAL};")
            else: self.score_lbl.setStyleSheet("color: red;")
            self.score_lbl.setText(f"Score: {self.score}"); self.next_round(); self.timer.start(3000)
        except: pass

class TrackingWidget(QWidget):
    def __init__(self):
        super().__init__()
        self.ball_pos = QPoint(200, 200)
        self.timer = QTimer(); self.timer.timeout.connect(self.move_ball)
        
    def start(self): self.timer.start(30)
    def stop(self): self.timer.stop()
    def move_ball(self):
        self.ball_pos += QPoint(random.randint(-5, 5), random.randint(-5, 5))
        self.ball_pos.setX(max(20, min(self.width()-20, self.ball_pos.x())))
        self.ball_pos.setY(max(20, min(self.height()-20, self.ball_pos.y())))
        self.update()
        
    def paintEvent(self, event):
        p = QPainter(self); p.fillRect(self.rect(), QColor("#000"))
        p.setBrush(QBrush(QColor(THEME_GOLD))); p.drawEllipse(self.ball_pos, 15, 15)
        p.setPen(QColor("#FFF")); p.drawText(20, 30, "FOLLOW THE ORB WITH YOUR EYES")

class MathReadingTask(QWidget):
    def __init__(self):
        super().__init__()
        layout = QVBoxLayout()
        
        mode_lay = QHBoxLayout()
        self.btn_track = QRadioButton("Visual Tracking"); self.btn_track.setChecked(True)
        self.btn_read = QRadioButton("Tech Reading")
        self.btn_track.toggled.connect(self.update_view)
        self.btn_read.toggled.connect(self.update_view)
        mode_lay.addStretch(); mode_lay.addWidget(self.btn_track); mode_lay.addWidget(self.btn_read); mode_lay.addStretch()
        layout.addLayout(mode_lay)

        self.stack = QStackedWidget()
        self.page_track = TrackingWidget()
        self.page_read = QWidget()
        l1 = QVBoxLayout()
        self.text_area = QTextEdit(); self.text_area.setReadOnly(True)
        self.text_area.setStyleSheet(f"font-size: 16px; color: {THEME_TEAL}; background: #111; padding: 20px;")
        l1.addWidget(QLabel("READ CAREFULLY:")); l1.addWidget(self.text_area)
        self.page_read.setLayout(l1)
        
        self.stack.addWidget(self.page_track); self.stack.addWidget(self.page_read)
        layout.addWidget(self.stack)
        self.setLayout(layout)
        self.articles = [
            "Neuroplasticity is the ability of neural networks in the brain to reorganize themselves by creating new neural connections throughout life. This allows neurons to compensate for injury and disease, and to adjust their activities in response to new situations. Researchers have discovered that neuroplasticity is not limited to childhood development but continues throughout adult life. This groundbreaking finding has revolutionized our understanding of brain function and has led to new therapeutic approaches for treating brain injuries, learning disabilities, and neurodegenerative diseases. The brain's remarkable ability to adapt and change forms the biological basis for learning new skills and forming new memories.",
            "Quantum entanglement is a phenomenon where two or more particles become interconnected in such a way that the quantum state of each particle cannot be described independently. When particles are entangled, they remain connected across vast distances, and measuring one particle instantaneously affects the state of the other. This counterintuitive phenomenon puzzled even Einstein, who called it 'spooky action at a distance.' Today, quantum entanglement is recognized as a fundamental aspect of quantum mechanics and has practical applications in quantum computing, quantum cryptography, and quantum teleportation. Scientists continue to explore the implications of entanglement for our understanding of reality and the nature of space and time.",
            "In cognitive science, attention is the cognitive process that allows us to focus on specific information while filtering out irrelevant stimuli. The human brain receives countless sensory inputs every second, yet we can only consciously process a fraction of this information. Selective attention mechanisms help us prioritize important information and maintain focus on relevant tasks. Research has shown that attention is not a single unified process but involves multiple neural systems and brain regions. Understanding attention mechanisms has profound implications for education, workplace productivity, mental health treatment, and the design of technology interfaces. Neuroscientists continue to investigate how attention shapes our perception and cognition."
        ]

    def start(self): self.update_view()
    def stop(self): self.page_track.stop()
    def start_track(self): 
        self.btn_track.setChecked(True)
        self.stack.setCurrentWidget(self.page_track)
        self.page_track.start()
    def start_reading(self): 
        self.btn_read.setChecked(True)
        self.stack.setCurrentWidget(self.page_read)
        self.page_track.stop()
        self.text_area.setText(random.choice(self.articles))
    def update_view(self):
        if self.btn_track.isChecked():
            self.stack.setCurrentWidget(self.page_track); self.page_track.start()
        else:
            self.stack.setCurrentWidget(self.page_read); self.page_track.stop(); self.text_area.setText(random.choice(self.articles))
    def get_score(self): return 2000

# =================================================================
# INTERACTIVE VISUALIZER
# =================================================================

class MindVisualizer(QWidget):
    def __init__(self):
        super().__init__()
        self.setSizePolicy(QSizePolicy.Policy.Expanding, QSizePolicy.Policy.Expanding)
        self.ball_y = 0.8; self.target_y = 0.8; self.jitter_intensity = 0.0
        self.ball_color = QColor(THEME_TEAL); self.state_label = "IDLE"
        self.timer = QTimer(); self.timer.timeout.connect(self.animate); self.timer.start(16)

    def update_data(self, label, s_rat, f_rat):
        self.state_label = label
        f_val = np.clip(f_rat, 0.5, 2.5); self.target_y = 1.0 - ((f_val - 0.5) / 2.0) 
        s_val = np.clip(s_rat, 0.5, 2.0); self.jitter_intensity = (s_val - 0.5) * 0.05 
        if label == "Stressed": self.ball_color = QColor(THEME_RED)
        elif label == "Focused": self.ball_color = QColor(THEME_GOLD)
        else: self.ball_color = QColor(THEME_TEAL)

    def animate(self):
        self.ball_y += (self.target_y - self.ball_y) * 0.05; self.update()

    def paintEvent(self, event):
        painter = QPainter(self); painter.setRenderHint(QPainter.RenderHint.Antialiasing)
        w, h = self.width(), self.height(); painter.fillRect(0, 0, w, h, QColor("#050508"))
        painter.setPen(QPen(QColor(30, 30, 40), 2))
        for i in range(0, w, 60): painter.drawLine(i, 0, i, h)
        for i in range(0, h, 60): painter.drawLine(0, i, w, i)
        
        jit_x = random.uniform(-self.jitter_intensity, self.jitter_intensity) * w
        jit_y = random.uniform(-self.jitter_intensity, self.jitter_intensity) * h
        cx = int(w * 0.5 + jit_x); cy = int(h * self.ball_y + jit_y); cy = max(50, min(h - 50, cy)); radius = 40
        
        glow = QRadialGradient(cx, cy, radius * 3)
        glow.setColorAt(0, QColor(self.ball_color.red(), self.ball_color.green(), self.ball_color.blue(), 100))
        glow.setColorAt(1, Qt.GlobalColor.transparent)
        painter.setBrush(QBrush(glow)); painter.setPen(Qt.PenStyle.NoPen)
        painter.drawEllipse(cx - radius*3, cy - radius*3, radius*6, radius*6)
        painter.setBrush(QBrush(self.ball_color)); painter.drawEllipse(cx - radius, cy - radius, radius*2, radius*2)
        
        painter.setPen(QPen(QColor("white"))); painter.setFont(QFont("Segoe UI", 16, QFont.Weight.Bold))
        painter.drawText(20, 40, f"STATUS: {self.state_label}")

# =================================================================
# MAIN PAGES
# =================================================================

class CalibrationPage(QWidget):
    def __init__(self, main_app):
        super().__init__()
        self.main_app = main_app
        layout = QVBoxLayout()
        layout.setContentsMargins(20, 20, 20, 20)
        layout.setSpacing(15)
        self.live_plot = pg.PlotWidget(title="Live EEG Signal Check")
        self.live_plot.setBackground(PANEL_BG); self.live_plot.setYRange(0, 4095); self.live_plot.setMinimumHeight(220)
        self.live_plot.setSizePolicy(QSizePolicy.Policy.Expanding, QSizePolicy.Policy.Expanding)
        self.plot_curve = self.live_plot.plot(pen=THEME_TEAL)
        self.plot_data = deque([0]*256, maxlen=256)
        layout.addWidget(self.live_plot)
        self.feedback = QLabel("Waiting for signal...")
        self.feedback.setStyleSheet(f"font-size: 16px; font-weight: bold; color: #888; border: 1px solid #444; padding: 12px; border-radius: 6px; background: #000;")
        self.feedback.setAlignment(Qt.AlignmentFlag.AlignCenter)
        self.feedback.setWordWrap(True)
        self.feedback.setMinimumHeight(60)
        layout.addWidget(self.feedback)
        status_row = QHBoxLayout()
        status_row.setSpacing(12)
        self.conn_lbl = QLabel("CONNECTION: DISCONNECTED")
        self.conn_lbl.setStyleSheet("color: #777; font-weight: bold;")
        self.conn_lbl.setMinimumWidth(220)
        self.strength_bar = QProgressBar()
        self.strength_bar.setRange(0, 100)
        self.strength_bar.setValue(0)
        self.strength_bar.setFormat("Signal: %p%")
        self.strength_bar.setTextVisible(True)
        self.strength_bar.setMinimumHeight(14)
        self.strength_bar.setStyleSheet(f"QProgressBar::chunk {{ background-color: {THEME_RED}; }}")
        status_row.addWidget(self.conn_lbl)
        status_row.addWidget(self.strength_bar, 1)
        layout.addLayout(status_row)
        self.task_stack = QStackedWidget()
        self.page_menu = QWidget(); menu_layout = QHBoxLayout()
        menu_layout.setContentsMargins(0, 0, 0, 0)
        menu_layout.setSpacing(15)
        self.card_base = self.create_task_card("BASELINE", "Relaxation", "Breathing Exercises", self.start_baseline)
        self.card_stress = self.create_task_card("STRESS", "High Load", "Math / Stroop", self.start_stress)
        self.card_focus = self.create_task_card("FOCUS", "Flow State", "Tracking / Reading", self.start_focus)
        menu_layout.addWidget(self.card_base); menu_layout.addWidget(self.card_stress); menu_layout.addWidget(self.card_focus)
        self.page_menu.setLayout(menu_layout)
        self.page_breath = BreathingTask(); self.page_stroop = StroopTask(); self.page_focus_task = MathReadingTask()
        self.task_stack.addWidget(self.page_menu); self.task_stack.addWidget(self.page_breath)
        self.task_stack.addWidget(self.page_stroop); self.task_stack.addWidget(self.page_focus_task)
        layout.addWidget(self.task_stack)
        ctrl = QFrame(); ctrl.setStyleSheet("background: #111; border: 1px solid #333; border-radius: 8px;")
        cl = QHBoxLayout()
        cl.setContentsMargins(15, 10, 15, 10)
        cl.setSpacing(15)
        self.status_lbl = QLabel("STATUS: IDLE", objectName="Subtitle")
        self.status_lbl.setStyleSheet("color: #AAA; font-size: 12px; letter-spacing: 1px;")
        self.xp_lbl = QLabel("NEURO XP: 0"); self.xp_lbl.setStyleSheet(f"font-size: 18px; font-weight: bold; color: {THEME_GOLD};")
        self.timer_lbl = QLabel("00:00"); self.timer_lbl.setStyleSheet(f"font-size: 24px; font-weight: bold; color: {THEME_TEAL};")
        self.btn_stop = QPushButton("ABORT SEQUENCE", objectName="DangerBtn"); self.btn_stop.setEnabled(False)
        self.btn_stop.setMinimumHeight(40)
        self.btn_stop.clicked.connect(self.stop_calibration)
        cl.addWidget(self.status_lbl); cl.addStretch(); cl.addWidget(self.xp_lbl); cl.addSpacing(20); cl.addWidget(self.timer_lbl); cl.addWidget(self.btn_stop)
        ctrl.setLayout(cl); layout.addWidget(ctrl)
        self.setLayout(layout); self.worker = None

    def create_task_card(self, title, subtitle, desc, func):
        f = QFrame(objectName="Card"); f.setMinimumHeight(200); f.setMinimumWidth(240)
        f.setSizePolicy(QSizePolicy.Policy.Expanding, QSizePolicy.Policy.Expanding)
        l = QVBoxLayout()
        l.setContentsMargins(15, 15, 15, 15)
        l.setSpacing(8)
        t = QLabel(title); t.setStyleSheet(f"font-size: 20px; font-weight: bold; color: {THEME_GOLD};"); t.setWordWrap(True)
        s = QLabel(subtitle); s.setStyleSheet("color: #AAA; font-style: italic;"); s.setWordWrap(True)
        d = QLabel(desc); d.setStyleSheet("color: #FFF;"); d.setWordWrap(True)
        b = QPushButton("INITIALIZE"); b.setCursor(Qt.CursorShape.PointingHandCursor); b.clicked.connect(func)
        b.setMinimumHeight(40)
        l.addWidget(t); l.addWidget(s); l.addWidget(d); l.addStretch(); l.addWidget(b)
        f.setLayout(l); return f

    def start_baseline(self):
        choice = self.ask_choice("Baseline Mode", "Choose breathing exercise:", [
            ("4-7-8", "4-7-8 (Calm)"),
            ("box", "Box (Focus)")
        ])
        if not choice:
            return
        self.page_breath.btn_478.setChecked(choice == "4-7-8")
        self.page_breath.btn_box.setChecked(choice == "box")
        self.setup_run("Baseline", self.page_breath)
        self.page_breath.start()

    def start_stress(self):
        choice = self.ask_choice("Stress Mode", "Choose stress task:", [
            ("stroop", "Stroop"),
            ("math", "Rapid Math")
        ])
        if not choice:
            return
        self.page_stroop.mode_stroop.setChecked(choice == "stroop")
        self.page_stroop.mode_math.setChecked(choice == "math")
        self.setup_run("Stressed", self.page_stroop)
        self.page_stroop.start()

    def start_focus(self):
        choice = self.ask_choice("Focus Mode", "Choose focus task:", [
            ("track", "Visual Tracking"),
            ("read", "Tech Reading")
        ])
        if not choice:
            return
        self.setup_run("Focused", self.page_focus_task)
        if choice == "track":
            self.page_focus_task.start_track()
        else:
            self.page_focus_task.start_reading()

    def ask_choice(self, title, text, options):
        msg = QMessageBox(self)
        msg.setWindowTitle(title)
        msg.setText(text)
        msg.setStyleSheet("background-color: #222; color: #FFF;")
        buttons = {}
        for key, label in options:
            buttons[key] = msg.addButton(label, QMessageBox.ButtonRole.ActionRole)
        msg.addButton("Cancel", QMessageBox.ButtonRole.RejectRole)
        msg.exec()
        clicked = msg.clickedButton()
        for key, btn in buttons.items():
            if clicked == btn:
                return key
        return None

    def setup_run(self, label, widget_page):
        port = self.main_app.get_port()
        if not port: return
        self.task_stack.setCurrentWidget(widget_page)
        self.worker = RecorderThread(port, label, duration=1200)
        self.update_connection("CONNECTING...", "#FF9800")
        self.update_strength(0)
        self.worker.progress_update.connect(self.update_progress)
        self.worker.live_signal.connect(self.update_plot)
        self.worker.feedback_msg.connect(self.update_feedback)
        self.worker.finished_phase.connect(self.phase_done)
        self.worker.error_occurred.connect(self.on_error)
        self.worker.signal_quality.connect(self.main_app.update_led)
        self.worker.connection_status.connect(self.update_connection)
        self.worker.signal_strength.connect(self.update_strength)
        self.worker.start()
        self.status_lbl.setText(f"RECORDING: {label.upper()}"); self.btn_stop.setEnabled(True)

    def stop_calibration(self):
        if self.worker: self.worker.stop(); self.worker.wait()
        self.phase_done("ABORTED")

    def phase_done(self, label):
        xp = 0
        if label != "ABORTED": xp = 2000
        if hasattr(self.task_stack.currentWidget(), 'get_score'): xp += self.task_stack.currentWidget().get_score()
        if hasattr(self.task_stack.currentWidget(), 'stop'): self.task_stack.currentWidget().stop()
        if xp > 0:
            f_path = CURRENT_USER_PATHS["HISTORY"]
            with open(f_path, 'a', newline='') as f:
                csv.writer(f).writerow([datetime.now().strftime("%Y-%m-%d %H:%M:%S"), f"SESSION COMPLETE: {label}", f"+{xp} XP"])
        QMessageBox.information(self, "MISSION COMPLETE", f"PROTOCOL FINISHED.\nTOTAL REWARD: {xp} NEURO XP")
        self.status_lbl.setText("SEQUENCE COMPLETE."); self.btn_stop.setEnabled(False)
        self.task_stack.setCurrentWidget(self.page_menu); self.xp_lbl.setText("NEURO XP: 0"); self.main_app.update_led(False)
        self.update_connection("DISCONNECTED", "#777")
        self.update_strength(0)

    def update_progress(self, pct, rem, count):
        mins, secs = divmod(rem, 60); self.timer_lbl.setText(f"{mins:02d}:{secs:02d}"); self.xp_lbl.setText(f"NEURO XP: {count // 25}")
    def update_plot(self, val):
        self.plot_data.append(val); self.plot_curve.setData(list(self.plot_data))
    def update_feedback(self, msg, color):
        self.feedback.setText(msg); self.feedback.setStyleSheet(f"font-size: 16px; font-weight: bold; color: {color}; border: 1px solid {color}; padding: 10px; border-radius: 6px; background: #000;")
    def update_connection(self, msg, color):
        self.conn_lbl.setText(f"CONNECTION: {msg}")
        self.conn_lbl.setStyleSheet(f"color: {color}; font-weight: bold;")
    def update_strength(self, val):
        self.strength_bar.setValue(val)
        if val >= 70:
            chunk = THEME_TEAL
        elif val >= 40:
            chunk = "#FF9800"
        else:
            chunk = THEME_RED
        self.strength_bar.setStyleSheet(f"QProgressBar::chunk {{ background-color: {chunk}; }}")
    def on_error(self, msg): QMessageBox.critical(self, "ERROR", msg); self.stop_calibration()

class TrainingPage(QWidget):
    def __init__(self):
        super().__init__()
        layout = QVBoxLayout()
        layout.setContentsMargins(20, 20, 20, 20)
        layout.setSpacing(15)
        title = QLabel("NEURAL NETWORK TRAINING", objectName="Title")
        title.setStyleSheet(f"font-size: 22px; font-weight: bold; color: {THEME_GOLD}; padding: 10px;")
        layout.addWidget(title)
        self.console = QTextEdit(); self.console.setReadOnly(True)
        self.console.setStyleSheet(f"background: #000; color: {THEME_TEAL}; font-family: monospace; border: 1px solid #333;")
        self.console.setMinimumHeight(260)
        self.console.setSizePolicy(QSizePolicy.Policy.Expanding, QSizePolicy.Policy.Expanding)
        layout.addWidget(self.console, 1)
        self.btn_demo = QPushButton("GENERATE DEMO DATA (TESTING)"); self.btn_demo.setCursor(Qt.CursorShape.PointingHandCursor)
        self.btn_demo.setStyleSheet(f"border: 1px dashed {THEME_GOLD}; color: {THEME_GOLD};")
        self.btn_demo.setMinimumHeight(44)
        self.btn_demo.clicked.connect(self.generate_demo)
        layout.addWidget(self.btn_demo)
        self.btn_train = QPushButton("EXECUTE TRAINING PIPELINE"); self.btn_train.setCursor(Qt.CursorShape.PointingHandCursor)
        self.btn_train.setMinimumHeight(44)
        self.btn_train.clicked.connect(self.start_training)
        layout.addWidget(self.btn_train)
        self.setLayout(layout); self.worker = None

    def generate_demo(self):
        self.console.clear(); self.console.append(">>> GENERATING SYNTHETIC DATASET...")
        QApplication.processEvents()
        try:
            count = generate_demo_dataset()
            self.console.append(f">>> SUCCESS. Generated {count} samples.\n>>> You can now EXECUTE TRAINING PIPELINE.")
        except Exception as e: self.console.append(f"ERROR: {str(e)}")

    def start_training(self):
        self.console.clear(); self.btn_train.setEnabled(False)
        self.worker = TrainerThread()
        self.worker.log_update.connect(lambda s: self.console.append(s))
        self.worker.training_finished.connect(lambda: self.btn_train.setEnabled(True))
        self.worker.start()

class MonitorPage(QWidget):
    def __init__(self, main_app):
        super().__init__()
        self.main_app = main_app
        layout = QVBoxLayout()
        layout.setContentsMargins(20, 20, 20, 20)
        layout.setSpacing(15)
        self.tabs = QTabWidget()
        self.tabs.setSizePolicy(QSizePolicy.Policy.Expanding, QSizePolicy.Policy.Expanding)
        self.tab_vis = QWidget(); vl = QVBoxLayout(); self.mind_vis = MindVisualizer()
        vl.setContentsMargins(0, 0, 0, 0)
        vl.setSpacing(10)
        vl.addWidget(self.mind_vis); self.tab_vis.setLayout(vl)
        self.tab_tech = QWidget(); tl = QGridLayout()
        tl.setContentsMargins(0, 0, 0, 0)
        tl.setSpacing(12)
        self.state_frame = QFrame(); self.state_frame.setStyleSheet(f"background: #080810; border: 2px solid {THEME_GOLD}; border-radius: 12px; padding: 10px;")
        self.state_frame.setMinimumHeight(160)
        sl = QVBoxLayout(); self.state_lbl = QLabel("IDLE"); self.state_lbl.setAlignment(Qt.AlignmentFlag.AlignCenter); self.state_lbl.setWordWrap(True)
        self.state_lbl.setStyleSheet(f"font-size: 72px; font-weight: bold; color: #333;")
        hl = QHBoxLayout(); self.tag_conf = QLabel("CONF: 0%"); self.tag_ver = QLabel("VER: ---")
        self.tag_conf.setWordWrap(True); self.tag_ver.setWordWrap(True)
        hl.addWidget(self.tag_conf); hl.addStretch(); hl.addWidget(self.tag_ver)
        sl.addWidget(self.state_lbl); sl.addLayout(hl); self.state_frame.setLayout(sl)
        self.graph = pg.PlotWidget(); self.graph.setBackground(BG_DARK); self.graph.showGrid(x=False, y=True, alpha=0.3)
        self.graph.setMinimumHeight(240)
        self.bars = pg.BarGraphItem(x=[1,2,3,4,5], height=[0,0,0,0,0], width=0.6, brush=THEME_TEAL, pen=None)
        self.graph.addItem(self.bars); self.graph.getAxis('bottom').setTicks([[(1,'D'),(2,'T'),(3,'A'),(4,'B'),(5,'G')]])
        tl.addWidget(self.state_frame, 0, 0); tl.addWidget(self.graph, 1, 0); self.tab_tech.setLayout(tl)
        self.tabs.addTab(self.tab_vis, "🚀 NEURO-GAME"); self.tabs.addTab(self.tab_tech, "📊 TECHNICAL DATA")
        layout.addWidget(self.tabs, 1)
        self.btn_start = QPushButton("INITIATE LIVE STREAM"); self.btn_start.setCursor(Qt.CursorShape.PointingHandCursor)
        self.btn_start.setMinimumHeight(44)
        self.btn_start.clicked.connect(self.toggle_monitor)
        layout.addWidget(self.btn_start)
        self.setLayout(layout); self.worker = None; self.monitoring = False

    def toggle_monitor(self):
        if not self.monitoring:
            port = self.main_app.get_port()
            if not port: return
            self.monitoring = True
            self.btn_start.setText("TERMINATE STREAM"); self.btn_start.setObjectName("DangerBtn"); self.btn_start.setStyleSheet("")
            self.worker = MonitorThread(port)
            self.worker.data_update.connect(self.update_ui)
            self.worker.dashboard_update.connect(self.main_app.page_dash.update_live_state)
            self.worker.signal_quality.connect(self.main_app.update_led)
            self.worker.error_occurred.connect(self.on_error)
            self.worker.start()
        else:
            self.monitoring = False
            self.btn_start.setText("INITIATE LIVE STREAM"); self.btn_start.setObjectName(""); self.btn_start.setStyleSheet("")
            if self.worker: self.worker.stop(); self.worker.wait(); self.reset_ui()

    def reset_ui(self):
        self.state_lbl.setText("IDLE"); self.state_lbl.setStyleSheet("color: #333; font-size: 72px;")
        self.main_app.page_dash.update_live_state("IDLE"); self.main_app.update_led(False); self.mind_vis.update_data("IDLE", 0, 0)
    def on_error(self, msg): QMessageBox.critical(self, "ERROR", msg); self.toggle_monitor()
    def update_ui(self, bands, label, conf, tag, s_rat, f_rat):
        self.bars.setOpts(height=bands); self.state_lbl.setText(label.upper())
        color = THEME_GOLD
        if label == "Stressed": color = THEME_RED
        elif label == "Focused": color = THEME_TEAL
        self.state_lbl.setStyleSheet(f"color: {color}; font-size: 72px; font-weight: bold;")
        self.tag_conf.setText(f"CONF: {conf*100:.1f}%"); self.tag_ver.setText(f"{tag} [S:{s_rat:.1f} F:{f_rat:.1f}]")
        self.mind_vis.update_data(label, s_rat, f_rat)

class DashboardPage(QWidget):
    def __init__(self, main_app=None):
        super().__init__()
        self.main_app = main_app
        main_layout = QVBoxLayout()
        main_layout.setContentsMargins(20, 20, 20, 20)
        main_layout.setSpacing(15)
        
        self.welcome_lbl = QLabel("MISSION DASHBOARD", objectName="Title")
        self.welcome_lbl.setStyleSheet(f"font-size: 22px; font-weight: bold; color: {THEME_GOLD}; margin: 0px; padding: 10px;")
        main_layout.addWidget(self.welcome_lbl)
        
        # Status Card
        self.state_card = QFrame()
        self.state_card.setStyleSheet(f"background: #080810; border-left: 6px solid #555; padding: 20px; min-height: 80px;")
        sl = QVBoxLayout()
        sl.setContentsMargins(10, 10, 10, 10)
        status_label = QLabel("LIVE STATUS")
        status_label.setStyleSheet(f"color: #AAA; font-size: 12px; text-transform: uppercase; letter-spacing: 1px;")
        sl.addWidget(status_label)
        self.lbl_s_val = QLabel("OFFLINE", objectName="StatVal")
        self.lbl_s_val.setStyleSheet(f"color: #AAA; font-size: 18px; font-weight: bold; min-height: 30px;")
        sl.addWidget(self.lbl_s_val)
        self.state_card.setLayout(sl)
        main_layout.addWidget(self.state_card)
        
        # Stats Cards
        gl = QGridLayout()
        gl.setHorizontalSpacing(15)
        gl.setVerticalSpacing(15)
        self.c_stress = self.make_card("STRESS THRESHOLD", "N/A", THEME_RED)
        self.c_focus = self.make_card("FOCUS THRESHOLD", "N/A", THEME_TEAL)
        self.c_data = self.make_card("NEURO XP", "0", THEME_GOLD)
        gl.addWidget(self.c_stress, 0, 0)
        gl.addWidget(self.c_focus, 0, 1)
        gl.addWidget(self.c_data, 0, 2)
        gl.setColumnStretch(0, 1)
        gl.setColumnStretch(1, 1)
        gl.setColumnStretch(2, 1)
        main_layout.addLayout(gl)
        
        # Event Log
        log_label = QLabel("EVENT LOG")
        log_label.setStyleSheet(f"color: #AAA; font-size: 12px; text-transform: uppercase; letter-spacing: 1px; margin: 10px 0px 5px 0px;")
        main_layout.addWidget(log_label)
        
        self.history = QListWidget(); self.history.setWordWrap(True)
        self.history.setStyleSheet("background: #000; border: 1px solid #333; color: #EEE; font-family: monospace; padding: 10px;")
        self.history.setMinimumHeight(200)
        main_layout.addWidget(self.history, 1)
        
        # Refresh Button
        btn = QPushButton("🔄 SYSTEM REFRESH")
        btn.setMinimumHeight(45)
        btn.setCursor(Qt.CursorShape.PointingHandCursor)
        if self.main_app:
            btn.clicked.connect(self.main_app.refresh_all)
        else:
            btn.clicked.connect(self.load_data)
        main_layout.addWidget(btn)
        
        self.setLayout(main_layout)
        self.load_data()

    def update_live_state(self, state):
        self.lbl_s_val.setText(state); c = "#555"
        if state == "STRESSED": c = THEME_RED
        elif state == "FOCUSED": c = THEME_TEAL
        elif state == "CALM": c = THEME_GOLD
        self.state_card.setStyleSheet(f"background: #080810; border-left: 6px solid {c};")

    def make_card(self, t, v, c):
        f = QFrame()
        f.setStyleSheet(f"background: #111; border-bottom: 3px solid {c}; padding: 20px; min-height: 120px;")
        f.setMinimumWidth(220)
        l = QVBoxLayout()
        l.setContentsMargins(10, 10, 10, 10)
        l.setSpacing(10)
        
        title_lbl = QLabel(t)
        title_lbl.setStyleSheet(f"color: #888; font-size: 11px; text-transform: uppercase; letter-spacing: 1px;")
        title_lbl.setWordWrap(True)
        l.addWidget(title_lbl)
        
        val = QLabel(v, objectName="StatVal")
        val.setStyleSheet(f"color: {c}; font-size: 24px; font-weight: bold; min-height: 35px;")
        val.setWordWrap(True)
        l.addWidget(val)
        l.addStretch()
        
        if "STRESS" in t: self.val_s = val
        elif "FOCUS" in t: self.val_f = val
        else: self.val_d = val
        
        f.setLayout(l)
        return f

    def load_data(self):
        if os.path.exists(CURRENT_USER_PATHS["PROFILE"]):
            try: self.welcome_lbl.setText(f"WELCOME BACK, {json.load(open(CURRENT_USER_PATHS['PROFILE'])).get('name', 'USER').upper()}")
            except: pass
        if os.path.exists(CURRENT_USER_PATHS["STATS"]):
            try:
                stats = pickle.load(open(CURRENT_USER_PATHS["STATS"], 'rb'))
                self.val_s.setText(f"{stats.get('stress_ratio', 0):.2f}"); self.val_f.setText(f"{stats.get('focus_ratio', 0):.2f}")
            except: pass
        if os.path.exists(CURRENT_USER_PATHS["RAW"]):
            try: self.val_d.setText(f"{len(pd.read_csv(CURRENT_USER_PATHS['RAW'], on_bad_lines='skip')):,}")
            except: pass
        self.history.clear()
        if os.path.exists(CURRENT_USER_PATHS["HISTORY"]):
            try:
                for row in list(csv.reader(open(CURRENT_USER_PATHS["HISTORY"])))[-15:]:
                    if len(row) > 1: self.history.addItem(f"[{row[0]}] {row[1]}")
            except: pass

class ProfilePage(QWidget):
    def __init__(self):
        super().__init__()
        main_layout = QVBoxLayout()
        main_layout.setContentsMargins(20, 20, 20, 20)
        main_layout.setSpacing(15)
        
        title = QLabel("USER PROFILE", objectName="Title")
        title.setStyleSheet(
            f"font-size: 22px; font-weight: bold; color: {THEME_GOLD}; margin: 0px; padding: 10px;"
            "background: qlineargradient(x1:0, y1:0, x2:1, y2:0, stop:0 #1A1A22, stop:1 #232331);"
            "border: 1px solid #2E2E3A; border-radius: 10px;"
        )
        main_layout.addWidget(title)
        
        form_frame = QFrame()
        form_frame.setStyleSheet(
            f"background: qlineargradient(x1:0, y1:0, x2:0, y2:1, stop:0 #14151A, stop:1 #0F1015);"
            f"border: 1px solid #2C2C36; border-radius: 12px; padding: 20px;"
        )
        form_layout = QFormLayout()
        form_layout.setContentsMargins(20, 20, 20, 20)
        form_layout.setSpacing(15)
        form_layout.setLabelAlignment(Qt.AlignmentFlag.AlignTop)
        form_layout.setRowWrapPolicy(QFormLayout.RowWrapPolicy.WrapAllRows)
        form_layout.setFormAlignment(Qt.AlignmentFlag.AlignTop)
        form_layout.setFieldGrowthPolicy(QFormLayout.FieldGrowthPolicy.ExpandingFieldsGrow)
        
        # Labels styling
        label_style = f"color: {THEME_TEAL}; font-weight: bold; min-height: 35px; letter-spacing: 0.5px;"
        
        name_label = QLabel("FULL NAME:")
        name_label.setStyleSheet(label_style)
        self.input_name = QLineEdit()
        self.input_name.setMinimumHeight(40)
        self.input_name.setStyleSheet("background: #0B0C10; border: 1px solid #2C3E50; color: #EDEDED; padding: 10px; border-radius: 8px;")
        self.input_name.setSizePolicy(QSizePolicy.Policy.Expanding, QSizePolicy.Policy.Fixed)
        form_layout.addRow(name_label, self.input_name)
        
        age_label = QLabel("AGE:")
        age_label.setStyleSheet(label_style)
        self.input_age = QLineEdit()
        self.input_age.setMinimumHeight(40)
        self.input_age.setStyleSheet("background: #0B0C10; border: 1px solid #2C3E50; color: #EDEDED; padding: 10px; border-radius: 8px;")
        self.input_age.setSizePolicy(QSizePolicy.Policy.Expanding, QSizePolicy.Policy.Fixed)
        form_layout.addRow(age_label, self.input_age)
        
        notes_label = QLabel("CLINICAL NOTES:")
        notes_label.setStyleSheet(label_style)
        self.input_notes = QTextEdit(); self.input_notes.setSizePolicy(QSizePolicy.Policy.Expanding, QSizePolicy.Policy.Expanding)
        self.input_notes.setStyleSheet("background: #0B0C10; border: 1px solid #3B2E5A; color: #EDEDED; padding: 10px; border-radius: 8px;")
        self.input_notes.setMinimumHeight(150)
        form_layout.addRow(notes_label, self.input_notes)
        
        form_frame.setLayout(form_layout)
        main_layout.addWidget(form_frame)
        
        btn = QPushButton("💾 SAVE PROFILE DATA")
        btn.setMinimumHeight(45)
        btn.setCursor(Qt.CursorShape.PointingHandCursor)
        btn.setStyleSheet(
            f"background: qlineargradient(x1:0, y1:0, x2:1, y2:0, stop:0 {THEME_GOLD}, stop:1 #F5D06F);"
            "color: #111; border: none; border-radius: 10px; font-weight: bold;"
        )
        btn.clicked.connect(self.save_profile)
        main_layout.addWidget(btn)
        
        main_layout.addStretch()
        self.setLayout(main_layout)
        self.load_profile()

    def save_profile(self):
        notes_text = self.input_notes.toPlainText().strip()
        existing = {}
        if os.path.exists(CURRENT_USER_PATHS["PROFILE"]):
            try:
                existing = json.load(open(CURRENT_USER_PATHS["PROFILE"], 'r'))
            except:
                existing = {}

        history = existing.get("notes_history", []) if isinstance(existing.get("notes_history", []), list) else []
        if notes_text:
            last_note = history[-1]["text"] if history else existing.get("notes", "")
            if notes_text != last_note:
                history.append({"timestamp": datetime.now().strftime("%Y-%m-%d %H:%M:%S"), "text": notes_text})

        data = {
            "name": self.input_name.text(),
            "age": self.input_age.text(),
            "notes": notes_text,
            "notes_history": history
        }
        try:
            with open(CURRENT_USER_PATHS["PROFILE"], 'w') as f: json.dump(data, f)
            QMessageBox.information(self, "SYSTEM", "PROFILE UPDATED.")
        except Exception as e: QMessageBox.critical(self, "ERROR", str(e))

    def load_profile(self):
        if os.path.exists(CURRENT_USER_PATHS["PROFILE"]):
            try:
                data = json.load(open(CURRENT_USER_PATHS["PROFILE"], 'r'))
                latest_notes = data.get("notes", "")
                if not latest_notes and isinstance(data.get("notes_history"), list) and data["notes_history"]:
                    latest_notes = data["notes_history"][-1].get("text", "")
                self.input_name.setText(data.get("name", "")); self.input_age.setText(data.get("age", "")); self.input_notes.setText(latest_notes)
            except: pass

class LoginPage(QDialog):
    def __init__(self):
        super().__init__()
        self.setWindowTitle("NEURO-MENTOR LOGIN")
        self.setStyleSheet(STYLESHEET)
        self.setModal(True)
        self.user = None

        screen = QApplication.primaryScreen()
        screen_size = screen.availableGeometry()
        width = min(720, int(screen_size.width() * 0.45))
        height = min(640, int(screen_size.height() * 0.55))
        self.resize(max(520, width), max(520, height))
        self.setMinimumSize(520, 520)

        layout = QVBoxLayout()
        layout.setSpacing(18)
        layout.setContentsMargins(30, 30, 30, 30)

        # Header
        title = QLabel("NEURO-MENTOR")
        title.setAlignment(Qt.AlignmentFlag.AlignCenter)
        title.setStyleSheet(f"font-size: 32px; font-weight: bold; color: {THEME_GOLD}; letter-spacing: 3px;")
        layout.addWidget(title)

        subtitle = QLabel("Multi-User Brain Computer Interface System")
        subtitle.setAlignment(Qt.AlignmentFlag.AlignCenter)
        subtitle.setStyleSheet("font-size: 12px; color: #AAA; font-style: italic;")
        layout.addWidget(subtitle)

        # Content Card
        card = QFrame()
        card.setStyleSheet("background: #111; border: 1px solid #333; border-radius: 10px;")
        card_layout = QVBoxLayout()
        card_layout.setContentsMargins(20, 20, 20, 20)
        card_layout.setSpacing(14)

        info_msg = QLabel("👤 Enter your username to login or create a new profile.\nEach user has isolated data and trained models.")
        info_msg.setWordWrap(True)
        info_msg.setStyleSheet("color: #AAA; font-size: 11px; line-height: 1.6;")
        card_layout.addWidget(info_msg)

        label = QLabel("USERNAME")
        label.setStyleSheet(f"color: {THEME_GOLD}; font-weight: bold; font-size: 11px; letter-spacing: 1px;")
        card_layout.addWidget(label)

        self.username = QLineEdit()
        self.username.setPlaceholderText("Enter your username (alphanumeric)")
        self.username.setMinimumHeight(40)
        self.username.setStyleSheet("padding: 10px 12px; border-radius: 6px;")
        self.username.returnPressed.connect(self.login)
        card_layout.addWidget(self.username)

        help_text = QLabel("💡 Use letters, numbers, and underscores only")
        help_text.setWordWrap(True)
        help_text.setStyleSheet("color: #666; font-size: 10px;")
        card_layout.addWidget(help_text)

        button_layout = QHBoxLayout()
        button_layout.setSpacing(10)

        btn_users = QPushButton("👥 VIEW PROFILES")
        btn_users.setMinimumHeight(44)
        btn_users.setStyleSheet(f"background: #222; border: 1px solid #444; color: {THEME_TEAL}; border-radius: 6px;")
        btn_users.clicked.connect(self.show_existing_users)
        button_layout.addWidget(btn_users)

        btn = QPushButton("🚀 ENTER SYSTEM")
        btn.setMinimumHeight(44)
        btn.clicked.connect(self.login)
        btn.setStyleSheet(f"background: {THEME_GOLD}; color: #000; border: none; border-radius: 6px; font-weight: bold; font-size: 14px;")
        btn.setCursor(Qt.CursorShape.PointingHandCursor)
        button_layout.addWidget(btn)

        card_layout.addLayout(button_layout)
        card.setLayout(card_layout)
        layout.addWidget(card)

        layout.addStretch()
        self.setLayout(layout)

    def show_existing_users(self):
        users_dir = os.path.join(BASE_DIR, "Resource files", "With LSTM", "profiles")
        if not os.path.exists(users_dir):
            QMessageBox.information(self, "PROFILES", "No profiles exist yet.")
            return
        
        users = [d for d in os.listdir(users_dir) if os.path.isdir(os.path.join(users_dir, d))]
        if not users:
            QMessageBox.information(self, "PROFILES", "No profiles exist yet.")
            return
        
        users_text = "\n".join([f"👤 {user}" for user in sorted(users)])
        QMessageBox.information(self, "EXISTING PROFILES", f"Available profiles:\n\n{users_text}")

    def login(self):
        username = self.username.text().strip()
        if not username:
            QMessageBox.warning(self, "Error", "Username cannot be empty")
            return
        
        # Validate username (alphanumeric and underscores only)
        if not all(c.isalnum() or c == '_' for c in username):
            QMessageBox.warning(self, "Error", "Username can only contain letters, numbers, and underscores")
            return
        
        # Create user profile directory and initialize if needed
        user_dir = os.path.join(BASE_DIR, "Resource files", "With LSTM", "profiles", username)
        if not os.path.exists(user_dir):
            try:
                os.makedirs(user_dir)
                self.initialize_user_profile(user_dir, username)
                QMessageBox.information(self, "NEW PROFILE", f"Welcome {username}!\nProfile created successfully.")
            except Exception as e:
                QMessageBox.critical(self, "Error", f"Failed to create profile: {str(e)}")
                return
        else:
            # Existing user - show welcome message
            profile_file = os.path.join(user_dir, "user_profile.json")
            user_name = username
            if os.path.exists(profile_file):
                try:
                    profile_data = json.load(open(profile_file))
                    user_name = profile_data.get("name", username)
                except:
                    pass
            # Uncomment the line below if you want a welcome message for returning users
            # QMessageBox.information(self, "WELCOME BACK", f"Welcome back, {user_name}!")
        
        self.user = username
        self.accept()

    def initialize_user_profile(self, user_dir, username):
        """Initialize a new user profile with default files"""
        # Create default profile.json
        profile_file = os.path.join(user_dir, "user_profile.json")
        default_profile = {
            "name": username,
            "age": "",
            "notes": "",
            "created_date": datetime.now().strftime("%Y-%m-%d %H:%M:%S"),
            "theme_preference": "dark"
        }
        with open(profile_file, 'w') as f:
            json.dump(default_profile, f, indent=2)
        
        # Create empty history CSV
        history_file = os.path.join(user_dir, "Session_History_Log.csv")
        with open(history_file, 'w', newline='') as f:
            csv.writer(f).writerow(["Timestamp", "Event", "Details"])
        
        # Create calibration data CSV
        raw_data_file = os.path.join(user_dir, "Raw_EEG_Training_Data.csv")
        with open(raw_data_file, 'w', newline='') as f:
            csv.writer(f).writerow(RAW_TRAINING_HEADERS)
        
        # Log initial setup
        with open(history_file, 'a', newline='') as f:
            csv.writer(f).writerow([datetime.now().strftime("%Y-%m-%d %H:%M:%S"), "PROFILE_CREATED", "New user profile initialized"])

class MainWindow(QMainWindow):
    def __init__(self, username):
        super().__init__()
        global CURRENT_USER_PATHS
        CURRENT_USER_PATHS = get_user_paths(username)

        self.setWindowTitle(f"NEURO-MENTOR // BCI SYSTEM [{username}]")
        # Responsive sizing based on screen
        screen = QApplication.primaryScreen()
        screen_size = screen.availableGeometry()
        width = max(1400, int(screen_size.width() * 0.95))
        height = max(900, int(screen_size.height() * 0.95))
        self.resize(width, height)
        self.setMinimumSize(1200, 800)
        self.setStyleSheet(STYLESHEET)
        central = QWidget()
        main = QHBoxLayout()
        main.setSpacing(0)
        main.setContentsMargins(0, 0, 0, 0)
        central.setLayout(main)
        self.setCentralWidget(central)
        
        sidebar = QFrame(objectName="NavPanel")
        sidebar.setMinimumWidth(240)
        sidebar.setMaximumWidth(300)
        sl = QVBoxLayout()
        sl.setSpacing(8)
        sl.setContentsMargins(15, 15, 15, 15)
        
        title = QLabel("NEURO\nMENTOR")
        title.setAlignment(Qt.AlignmentFlag.AlignCenter)
        title.setStyleSheet(f"font-size: 28px; font-weight: bold; color: {THEME_GOLD}; letter-spacing: 3px; margin-bottom: 15px; min-height: 80px;")
        title.setWordWrap(True)
        sl.addWidget(title)
        
        self.stack = QStackedWidget(); self.nav_btns = []
        
        # Instantiate Pages Explicitly
        self.page_dash = DashboardPage(self)
        self.page_prof = ProfilePage()
        self.page_cal = CalibrationPage(self)
        self.page_train = TrainingPage()
        self.page_mon = MonitorPage(self)

        self.page_dash_view = self.wrap_scroll(self.page_dash)
        self.page_prof_view = self.wrap_scroll(self.page_prof)
        self.page_cal_view = self.wrap_scroll(self.page_cal)
        self.page_train_view = self.wrap_scroll(self.page_train)
        self.page_mon_view = self.wrap_scroll(self.page_mon)
        
        pages = [("DASHBOARD", self.page_dash_view), ("PROFILE", self.page_prof_view), ("CALIBRATE", self.page_cal_view), 
             ("NEURAL NET", self.page_train_view), ("LIVE FEED", self.page_mon_view)]
        
        for idx, (name, page_widget) in enumerate(pages):
            btn = QPushButton(name)
            btn.setObjectName("MenuBtn")
            btn.setCheckable(True)
            btn.setCursor(Qt.CursorShape.PointingHandCursor)
            btn.setMinimumHeight(40)
            btn.clicked.connect(lambda _, i=idx: self.switch_page(i))
            sl.addWidget(btn)
            self.nav_btns.append(btn)
            self.stack.addWidget(page_widget)
        
        sl.addStretch()
        
        # Signal indicator with label
        signal_frame = QFrame()
        signal_frame.setStyleSheet(f"background: transparent; border: none;")
        led_lay = QHBoxLayout()
        led_lay.setContentsMargins(0, 0, 0, 0)
        led_lay.setSpacing(8)
        signal_lbl = QLabel("SIGNAL:")
        signal_lbl.setStyleSheet(f"color: #888; font-size: 11px; min-width: 50px;")
        self.led = QLabel()
        self.led.setFixedSize(16, 16)
        self.led.setStyleSheet("background: #333; border-radius: 8px;")
        self.signal_status_lbl = QLabel("IDLE")
        self.signal_status_lbl.setStyleSheet("color: #777; font-size: 11px;")
        self.signal_status_lbl.setMinimumWidth(70)
        led_lay.addWidget(signal_lbl)
        led_lay.addWidget(self.led)
        led_lay.addWidget(self.signal_status_lbl)
        signal_frame.setLayout(led_lay)
        sl.addWidget(signal_frame)
        
        # COM Port selector
        com_label = QLabel("COM PORT:")
        com_label.setStyleSheet(f"color: #888; font-size: 11px;")
        sl.addWidget(com_label)
        self.com_box = QComboBox()
        self.com_box.setMinimumHeight(35)
        self.refresh_ports()
        sl.addWidget(self.com_box)
        
        # Switch User Button
        btn_switch = QPushButton("🔄 SWITCH USER")
        btn_switch.setMinimumHeight(40)
        btn_switch.setStyleSheet(f"background: #222; border: 1px solid #555; color: {THEME_TEAL}; border-radius: 4px;")
        btn_switch.setCursor(Qt.CursorShape.PointingHandCursor)
        btn_switch.clicked.connect(self.switch_user)
        sl.addWidget(btn_switch)
        
        sidebar.setLayout(sl)
        main.addWidget(sidebar)
        self.stack.setSizePolicy(QSizePolicy.Policy.Expanding, QSizePolicy.Policy.Expanding)
        main.addWidget(self.stack)
        self.nav_btns[0].setChecked(True)

    def refresh_ports(self):
        self.com_box.clear()
        for p in serial.tools.list_ports.comports():
            self.com_box.addItem(p.device)

    def refresh_all(self):
        if (self.page_cal.worker and self.page_cal.worker.isRunning()) or self.page_mon.monitoring:
            QMessageBox.information(self, "SYSTEM REFRESH", "Refresh will update UI and ports without interrupting active sessions.")

        current = self.com_box.currentText()
        self.refresh_ports()
        if current:
            idx = self.com_box.findText(current)
            if idx >= 0:
                self.com_box.setCurrentIndex(idx)

        self.page_dash.load_data()
        self.page_prof.load_profile()
        self.update_led(False)

    def get_port(self):
        p = self.com_box.currentText()
        if not p: QMessageBox.critical(self, "ERROR", "SELECT COM PORT"); return None
        return p

    def switch_page(self, i):
        self.stack.setCurrentIndex(i)
        for b in self.nav_btns: b.setChecked(False)
        self.nav_btns[i].setChecked(True)
        if i == 0: self.page_dash.load_data()

    def wrap_scroll(self, widget):
        area = QScrollArea()
        area.setWidgetResizable(True)
        area.setFrameShape(QFrame.Shape.NoFrame)
        area.setHorizontalScrollBarPolicy(Qt.ScrollBarPolicy.ScrollBarAsNeeded)
        area.setVerticalScrollBarPolicy(Qt.ScrollBarPolicy.ScrollBarAsNeeded)
        area.setWidget(widget)
        return area

    def switch_user(self):
        reply = QMessageBox.question(self, "SWITCH USER", 
            "This will end the current session and return to login. Continue?",
            QMessageBox.StandardButton.Yes | QMessageBox.StandardButton.No)
        if reply != QMessageBox.StandardButton.Yes:
            return

        # Prevent switching during active training
        if self.page_train.worker and self.page_train.worker.isRunning():
            QMessageBox.warning(self, "SWITCH USER", "Training is in progress. Please wait for it to finish before switching users.")
            return

        # Stop live sessions safely
        if self.page_mon.monitoring:
            self.page_mon.toggle_monitor()
        if self.page_cal.worker:
            self.page_cal.stop_calibration()

        login = LoginPage()
        if login.exec() != QDialog.DialogCode.Accepted:
            return

        username = login.user
        global CURRENT_USER_PATHS
        CURRENT_USER_PATHS = get_user_paths(username)
        self.setWindowTitle(f"NEURO-MENTOR // BCI SYSTEM [{username}]")

        # Reset/refresh UI for new user
        self.page_dash.load_data()
        self.page_prof.load_profile()
        self.page_train.console.clear()
        self.page_cal.feedback.setText("Waiting for signal...")
        self.page_cal.task_stack.setCurrentWidget(self.page_cal.page_menu)
        self.page_cal.status_lbl.setText("STATUS: IDLE")
        self.page_cal.xp_lbl.setText("NEURO XP: 0")
        self.page_cal.timer_lbl.setText("00:00")
        self.page_cal.btn_stop.setEnabled(False)
        self.page_cal.update_connection("DISCONNECTED", "#777")
        self.page_cal.update_strength(0)
        self.page_mon.reset_ui()
        self.update_led(False)

    def update_led(self, good):
        col = THEME_TEAL if good else "#333"
        glow = f"box-shadow: 0 0 8px {col};" if good else ""
        self.led.setStyleSheet(f"background: {col}; border-radius: 6px; {glow}")
        if good:
            self.signal_status_lbl.setText("GOOD")
            self.signal_status_lbl.setStyleSheet(f"color: {THEME_TEAL}; font-size: 11px;")
            self.led.setToolTip("Signal quality: GOOD")
        else:
            self.signal_status_lbl.setText("NO SIGNAL")
            self.signal_status_lbl.setStyleSheet("color: #777; font-size: 11px;")
            self.led.setToolTip("Signal quality: NO SIGNAL")

if __name__ == "__main__":
    app = QApplication(sys.argv)
    login = LoginPage()
    if login.exec() == QDialog.DialogCode.Accepted:
        window = MainWindow(login.user)
        window.show()
        sys.exit(app.exec())
```
